# Optic nerve innervation promotes Wnt/b-catenin pathway activity and progenitor cell proliferation in the zebrafish optic tectum

**DOI:** 10.64898/2026.06.17.732896

**Authors:** Olivia Hagen, Yehyun Kim, Elaine Kushkowski, Jiahe Yue, Hannah Rouse, Sara Helmstetter, Cameron Roberts, Máté Varga, Steve W. Wilson, Kara L. Cerveny

## Abstract

In the zebrafish visual system, accurate retinotopic mapping occurs throughout life as new neurons are generated and integrated into existing circuitry in both the retina and optic tectum (OT). To explore how OT development changes relative to innervation from the retina, we examined cell death and proliferation in genetic and surgical models that disrupt retinal innervation of the OT. Specifically, we compared *lakritz (lak)* mutants, which have no optic nerves due to a lesion in the *atoh7* gene, with either wild-type or one-eyed fish generated through surgical eye removal. We observed elevated cell death, fewer proliferating progenitors, and fewer *sox2+* OT neuroepithelial stem cells in *lak* mutant and denervated OT lobes. To examine whether light-mediated vision contributes to proliferation and survival in the optic tectum, we reared fish in constant darkness and then compared survival and proliferation of OT cells in innervated and non-innervated tecta. We found that OT cells were still more likely to survive and proliferate in the presence of optic nerve innervation even when fish were reared in the dark. To identify molecular pathways that could regulate OT growth, we examined the expression of known mitogens in the zebrafish optic tectum and found evidence that Wnt/β-catenin pathway activity could promote innervation-dependent proliferation in *lak* mutant tecta. Expression of both *wnt3a* and the Wnt/β-catenin target gene *axin2,* as detected by *in situ* hybridization and RT-qPCR, is decreased in non-innervated tectal lobes. Further supporting an innervation-dependent role for Wnt/β-catenin pathway activation in the zebrafish OT, we found that *lak* mutants treated with a Wnt-pathway agonist, BIO, exhibited levels of OT cell proliferation that were indistinguishable from wild-type. Together these findings suggest that progenitor cells in the optic tectum produce Wnt3a in response to innervation by the optic nerve, providing new insight into how a vertebrate visual system coordinates growth across its sensory and recipient tissues.

## Introduction

Lifelong neurogenesis is observed in all vertebrates where it undergirds learning and memory (reviewed in (Chen et al., 2025; Gonçalves et al., 2016; Konefal et al., 2013; Miller and Sahay, 2019)) as well as repair and regeneration (reviewed in (Alunni and Bally-Cuif, 2016; Becker and Becker, 2022; Lindsey et al., 2018). In teleost fishes such as zebrafish (*Danio rerio*), life-long growth of the nervous system is supported primarily by dedicated populations of stem and progenitor cells (Foley et al., 2024) that are located in neuroepithelial stem cell niches. Specifically in the visual system, growth is fueled by two stem cell niches: the ciliary marginal zone (CMZ) in the retina and the tectal proliferation zone (TPZ) in the optic tectum (OT) (Cerveny et al., 2012; Raymond and Easter, 1983; Raymond et al., 1983; Sato et al., 2016; Sokolova et al., 2023). Progenitors within the CMZ generate new neurons around the most peripheral edge of the functional retina (e.g., (Tsingos et al., 2019; Wan et al., 2016). Experiments in fishes and amphibians provide data that gene expression programs guiding retinal development are recapitulated for life-long growth within the CMZ (Perron et al., 1998; Xu et al., 2020). Proliferating progenitors within the TPZ add concentric crescents of new neurons to the medial and caudal edges of the optic tectum (Recher et al., 2013; Sato et al., 2015). Within both the CMZ and TPZ new neurons seem to be pushed out of the stem cell niche as progenitors proliferate and then differentiate.

The molecular mechanisms that coordinate cell behaviors within the CMZ and TPZ are ripe for active investigation. Historically, many studies have connected retinal input to optic tectum growth as atrophy of the tectum has been observed upon elimination of retinal input in various species of fish at larval and adult stages (e.g. Currie and Cowan, 1974; Kollros, 1982; Larsell, 1931; Raymond et al., 1983; Schmatolla, 1972; Schmatolla and Erdmann, 1973; White, 1948). Innervation of the optic tectum by retinal ganglion cell (RGC) axons appears to precede much of the cell proliferation and differentiation in the teleost optic tectum, coupling visual system development and growth. For instance, Raymond et al., (1983) showed that in adult goldfish, permanent removal of optic input by unilateral eye extirpation led to a sustained depression of ^3^H-thymidine incorporation on the denervated side of the OT. When optic input was temporarily disrupted by optic nerve crush, the number of ^3^H-thymidine-labelled cells initially decreased but then increased dramatically as new optic nerve fibers migrated into the denervated OT.

Despite a wealth of studies examining OT growth and development, the cellular and molecular cues that contribute to innervation-dependent growth are only starting to be elucidated. A number of extrinsically-regulated pathways involved in regional specific growth and neurogenesis have been implicated in OT growth control in adult fishes, and these pathways may also be important for innervation-dependent growth and development during embryonic and larval periods. For example, the Delta-Notch pathway modulates the number of new neurons generated in the adult OT, with Notch activation dampening proliferation of adult OT stem cells (Dozawa et al., 2014). The extent of Notch pathway activity may balance persistent *her5+* neuroepithelial cells that proliferate and give rise to *her4+* radial glial cells, which, in turn, divide to generate new neurons (Galant et al., 2016). In addition, both the Shh and Wnt/β-catenin pathways influence stem and progenitor cell behaviors in the adult zebrafish OT. Shh pathway activity correlates with maintenance of *sox2+* neural stem cells and Wnt/β-catenin pathway activity promotes proliferation over cell cycle exit and differentiation (Shitasako et al., 2017).

To begin to investigate the cellular and molecular control of innervation-dependent growth and development of the zebrafish OT, we compared cell behaviors in wild-type larvae, *lakritz (lak)* mutant larvae that lack retinal ganglion cells and therefore have no optic nerves, and one-eyed wild-type larvae (Fig 1). With a combination of cellular and pharmacological approaches, our data reveal a critical period during which survival, proliferation, and differentiation of OT cells require optic nerve innervation and highlight an instructive role for innervation in activating Wnt/β-catenin pathway activity to promote optic-nerve dependent proliferation in the OT.

**Figure 1:**
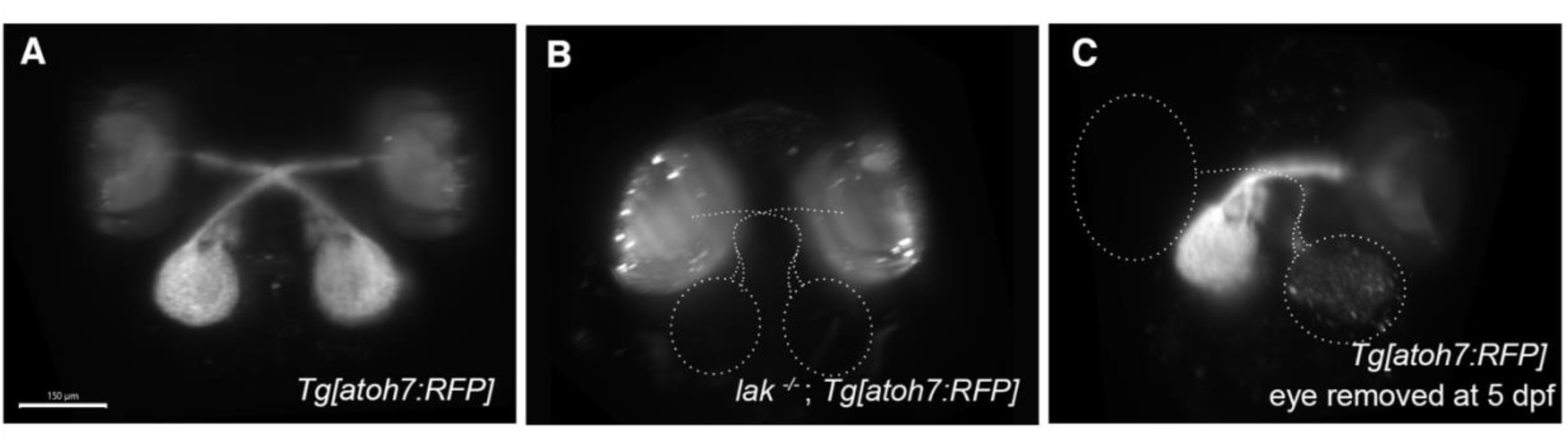
Examples of larvae used for assessing the influence of optic nerve innervation on tectal growth and development. **(A-C)** Maximum intensity projections of membrane-targeted RFP in *atoh7*-expressing retinal cells of living 9 dpf larvae captured with selective plane illumination microscopy demonstrating (A) typical innervation of the tectum by the optic nerve, visible as the axons from the *atoh7+* retinal ganglion cells, (B) absence of optic nerve and tectal innervation in *lak^-/-^* mutants (only retinae are apparent by low levels of RFP-expression), and (C) loss of optic nerve and tectal innervation when the left eye is removed at 5 dpf. Dotted lines indicate the absent eye in C and non-innervated tecta in B and C. Bar is 150 µm.

## Results

### Optic tectum cells require optic nerve input for survival

Cell-cell connections from the eye to the brain have been linked to neuron survival in the optic tectum. For example, studies of adult cave fish (Soares et al., 2004) and one-eyed larval *Fundulus* fish (White, 1948) show that connections between the retina and optic tectum (OT) contribute to tectal size and complexity. To investigate how connections from the retina to the OT influence cell survival, we documented the number of TUNEL+ cells at distinct times in the OT of *lakritz* (*lak*) mutants, which lack retinal ganglion cells, which form the optic nerve (Kay et al., 2001). We found that cell survival in the zebrafish OT is correlated with innervation from the retina beginning at 7 dpf (Fig 2A-I), approximately 4 days after most RGC axons have connected with their tectal partners (Burrill and Easter, 1994; Stuermer, 1988). Similar numbers of TUNEL+ cells were observed in wild-type and *lak* mutant tecta at both 5 and 6 dpf (Figure 2A-D; 2I). By 7 dpf, *lak* tecta contained nearly twice TUNEL+ cells. Two days later at 9 dpf, both wild-type and *lak* brains contained a similar, elevated number of TUNEL+ cells. Statistical analysis indicates of a strong contribution of both genotype and age as the average number of TUNEL+ cells relative to genotype and age appears distinctly different, with both variables exhibiting an interaction effect (ANOVA, f(3)=6.43, p<0.001). Pairwise comparisons of *lak* and wild-type data at each age showed that the average numbers of TUNEL+ cells were higher in *lak* mutants compared to wild-type at 7 dpf but not at other time points (Wilcoxon SRT, p<0.001). In addition, the average number of TUNEL+ cells was larger in wild-type OT at 9 dpf relative to all earlier wild-type stages (Tukey HSD p <0.002 for each stage comparison). See Table S3-S4 for results of statistical analyses. Taken together these data provide support for specifically-timed developmental apoptosis in the growing OT of zebrafish larvae that is accelerated in the absence of innervation.

**Figure 2:**
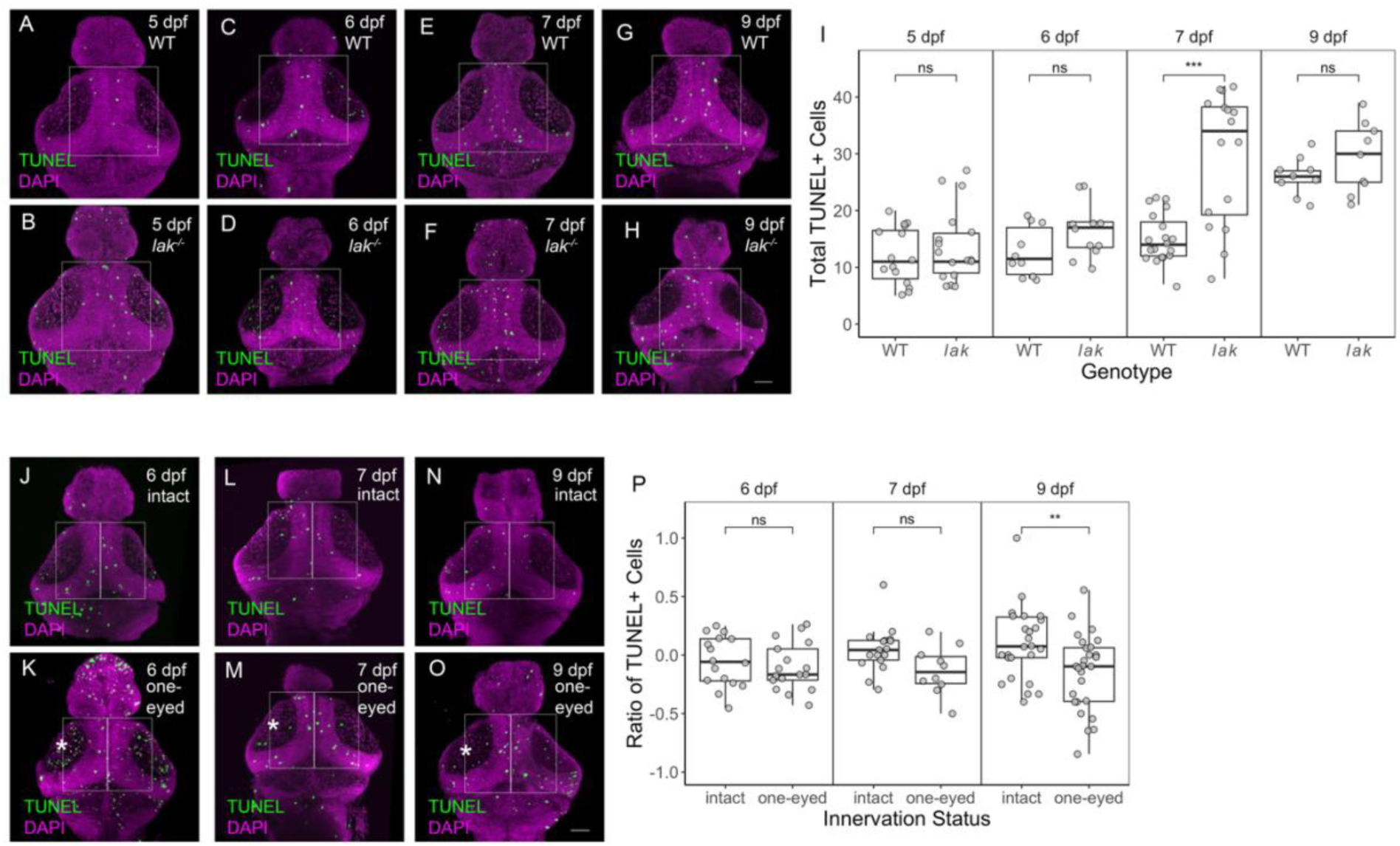
Tectal cell apotosis increases in the OT when optic nerve innervation is absent or lost. **(A-H)** Maximum intensity projections of dorsal views of dissected brains from wild-type (A, C, E, G) and *lak^-/-^* (B, D, F, H) larvae showing TUNEL+ cells (green) and nuclei (DAPI, magenta). Bar is 50 µm. Ages and genotypes indicated in the upper right corner of all panels. TUNEL+ cells in the boxed region of the optic tectum, excluding those in the tectal neuropil, were counted and compared. **(I)** Box plots overlaid with individual data points show TUNEL+ cells as a function of genotype at ages indicated. Abbreviations: ns, not significant; *** p ≤ 0.001 (Wilcoxon ranked sums test). **(J-O)** Maximum intensity projections of dorsal views of dissected brains from age-matched wild-type intact (J, L, N) and one-eyed (K, M, O) larvae showing TUNEL+ cells (green) and nuclei (DAPI, magenta). Bar is 50 µm. Ages indicated in the upper right corner of all panels. Innervated tectal lobes indicated with an asterisk. TUNEL+ cells inside boxed regions, excluding those in the tectal neuropil, were counted and compared. **(P)** Box plots overlaid with individual data points display TUNEL+ cells as a ratio between the left (or innervated post-extirpation) and right (or denervated post-extirpation) sides divided by the total TUNEL+ cells in the boxed regions of the optic tectum at ages indicated. n ≥ 10 for each condition. Abbreviations: ns, not significant; ** p ≤ 0.01 (Wilcoxon ranked sums test).

To further examine the connection between innervation and tectal cell survival, we compared the number of TUNEL+ cells in optic tecta from larvae subjected to unilateral eye extirpation at 5 dpf to intact (two-eyed) larvae. These comparisons revealed a requirement for innervation-dependent cell survival 4 days after eye removal (9 dpf). Visual inspection of the image data suggested that the number of TUNEL+ cells appeared to increase dramatically throughout the OT of one-eyed fish soon after eye-loss, especially in the neuropil. Although most of our quantitative analyses excluded the tectal neuropil, focusing only on cells in and adjacent to the TPZ, we observed elevated numbers of TUNEL+ spots within the neuropil regions of the OT, on both sides of the brain one day after eye extirpation (Fig S1A-B).Focusing only on the TPZ and tectal neurons revealed increased numbers of TUNEL+ apoptotic cells several days after retinal injury/loss of innervation (Fig 2J-P). The numbers of TUNEL+ cells around the midline of each side of the tectum (e.g., boxes in Fig 2J-O) were counted and compared by subtracting the number of TUNEL+ cells in the innervated (left) side from the denervated (right) side and then dividing the difference by the total number of TUNEL+ cells. Indicating that both sides of the optic tectum exhibit similar amounts of apoptosis, the average difference ratio of TUNEL+ cells in tecta from two-eyed larvae was close to zero for all ages examined (Fig 2P). Statistical analysis revealed that innervation status contributed significantly to differences between intact and one-eyed larvae (Fig 2J-P; ANOVA, f(1)=12.48, p<0.001). Pairwise comparisons showed significantly more TUNEL+ cells in the denervated tectal lobe only at 9 dpf (Wilcoxon SRT, p<0.007). As above, see Tables S2-S4 for statistics.

Together the data from our investigations of cell death identify a time-dependent requirement for innervation-dependent cell survival, with more tectal cells dying in non-innervated tectal lobes (*lak* mutants) or after loss of innervation (one-eyed larvae).

### Optic tectum cells require optic nerve input for proliferation

Tissue size depends on both cell survival and cell proliferation, with life-long growth requiring sustained and carefully controlled proliferation from a population of stem and progenitor cells. We initially analyzed tectal cell proliferation between 5-9 dpf and found that the amount of proliferation in the OT depends on innervation beginning around 9 dpf (Fig 3). Actively proliferating cells were labeled for approximately 16 hours with EdU before fixation at either 5, 7, or 9 dpf. Wild-type and *lak* mutant tecta contained similar numbers of EdU+ cells (Fig 3A-B, G) at 5 dpf. By 7 dpf, the average number of EdU+ cells in both wild-type and *lak* tecta noticeably decreased (Fig 3C-D, G). By 9 dpf, the average number of EdU+ cells in wild-type tecta was similar to the numbers of proliferating cells in wild-type and *lak* tecta at 7 dpf (Fig 3E-F, G). However, the average number of EdU+ cells in *lak* mutant tecta at 9 dpf decreased nearly 3-fold. Statistical analyses suggest that both age and genotype influence proliferation (ANOVA, f(2)=3.215, p<0.05) with age as the main driver of the observed differences (ANOVA, f(2)=43.05, p<0.001). See Tables S2-S4 for all results of our statistical analyses.

**Figure 3:**
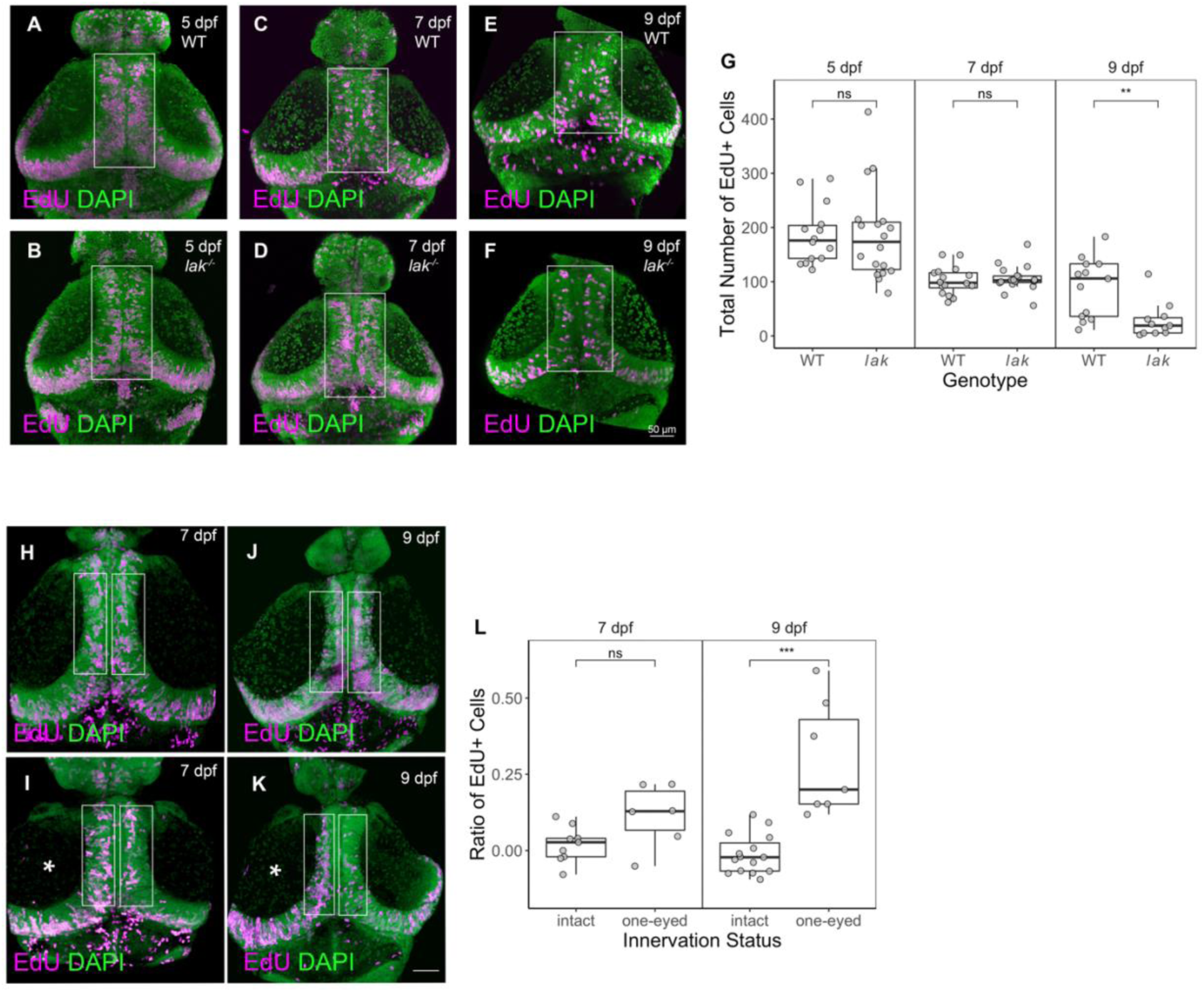
The extent of cell proliferation in the tectum is dependent on retinal innervation after 7 dpf. **(A-H)** Maximum intensity projections of dorsal views of dissected brains from wild-type (A, C, E) and *lak^-/-^* (B, D, F) larvae showing EdU+ cells (magenta) and nuclei (DAPI, green). Bar is 50 µm. Ages and genotypes indicated in upper right corner of all panels. EdU+ cells in the boxed regions in the optic tectum, excluding those in the neuropil, were counted and compared. **(G)** Box plots overlaid with individual data points show EdU+ cells as a function of genotype and age. n ≥ 11 for each condition. Abbreviations: ns, not significant; ** p ≤ 0.01 (Wilcoxon ranked sums test). **(H-K)** Maximum intensity projections of dorsal views of dissected brains from wild-type intact (H, J) and one-eyed (I, K) larvae showing EdU+ cells (magenta) and nuclei (DAPI, green). Bar is 50 µm. Ages indicated in upper right corner of all panels. Innervated tectal lobes indicated with an asterisk. EdU+ cells inside boxed regions, excluding those in the neuropil, were counted and compared. **(L)** Box plots overlaid with individual data points show EdU+ cells as a ratio between the left (innervated post-extirpation) and right (denervated post-extirpation) sides of the optic tectum at ages indicated. n ≥ 6 for each condition. Abbreviations: ns, not significant; * p ≤ 0.05, *** p ≤ 0.001 (Wilcoxon ranked sums test).

Despite fewer EdU+ cells in *lak* mutants, the rate of tectal neuron differentiation appears similar in wild-type and *lak* mutants. When larvae are pulsed with EdU for 4 hours at 8 dpf and then examined for HuC/elav3 expression 2 days or 4 days later, fewer double-positive neurons are observed in *lak* mutants at all timepoints (Fig S2A-C). This initially suggested delayed differentiation in *lak* mutants. However, linear regression analysis of the rate at which EdU+ cells are chased into HuC/elav3+ cells suggests that the differentiation rate is nearly identical regardless of genotype (Fig S2D-E). Taken together with the analysis of rate of change of EdU+ cells (Fig S2F), these pulse-chase data are consistent with elevated levels of apoptosis (e.g., Fig 1), decreased probability of entering the cell cycle and/or a slower cell cycle (e.g.,(Sherman et al., 2023)) contributing to fewer EdU+ tectal stem cells and thus fewer EdU+ neurons in *lak* mutants. Although *lak* mutants exhibit elevated apoptosis and decreased proliferation, overall brain size, does not appear noticeably smaller at these early larval stages (Fig S3, Tables S3 and S4).

We also compared the number of EdU+ cells in tecta from wild-type two-eyed larvae and larvae subjected to unilateral eye extirpation at 5 dpf. These data indicate that significant\ differences in proliferation emerge several days after injury/loss of retinal input (Fig 3H-L). The numbers of EdU+ cells on each side of the midline (e.g., boxes in Fig 3H-K) were counted and compared by subtracting the number of EdU+ cells in the innervated (left) side from the denervated (right) side and then dividing that difference by the total number of EdU+ cells. Indicating that cells on both sides of tectum proliferated at the same rate, we found that the average difference ratio (L-R/total) of EdU+ cells in tecta from two-eyed larvae was close to zero for the ages examined (Fig 3L). Statistical analysis revealed that innervation status contributed significantly to differences between intact and one-eyed larvae (Fig 3H-L; ANOVA, f(1)=35.963, p<0.001), with an interaction effect detected between age and innervation status (ANOVA, f(1)=8.868, p=0.006). Pairwise comparisons showed significantly fewer EdU+ cells in the denervated tectal lobe only by 9 dpf (Tukey HSD, p=0.001).

Together, these EdU incorporation data are consistent with OT stem and progenitor cells proliferating regardless of innervation status for the first ∼6 dpf. After 7 dpf, stem and progenitor cells in the OT appear to modulate their proliferation in response to innervation, with *lak* mutants showing dramatically decreased proliferation after 7 dpf and denervated brains showing many fewer proliferating cells 4 days after eye removal.

### Physical connections between the eye and brain via the optic nerve link optic tectum cell survival and proliferation

Although blocking RGC activity during early stages of visual system development does not alter retinotopic mapping of RGC axons in the OT (Stuermer et al., 1990), recent studies suggest that fish reared under low light conditions when RGCs are less active (Hall and Tropepe, 2018) or without optic nerve innervation due to the *lakritz* mutation (Figure 3, this study; Sherman et al., 2023) exhibit decreased proliferation in the OT. To examine whether neuronal activity through the optic nerve input might promote both survival and cell proliferation in the OT, we reared intact and one-eyed fish in a nearly constant dark environment and compared them with intact and one-eyed fish reared in a typical light-dark environment with visual stimulation from conspecifics and other movement in the room. The difference ratio of dying or proliferating cells on the innervated and denervated sides was calculated and compared both within and across environmental light conditions (representative images in Fig 4A-D for cell survival and Fig 4F-I for cell proliferation). For both cell death and cell proliferation, innervation status was the primary driver of all differences observed (Fig 4E and J) with environmental light conditions having a small, non-statistically significant contribution. The total number of cells proliferating or dying in intact wild-type larvae did not vary between light-dark and dark conditions (Fig S4).

**Figure 4:**
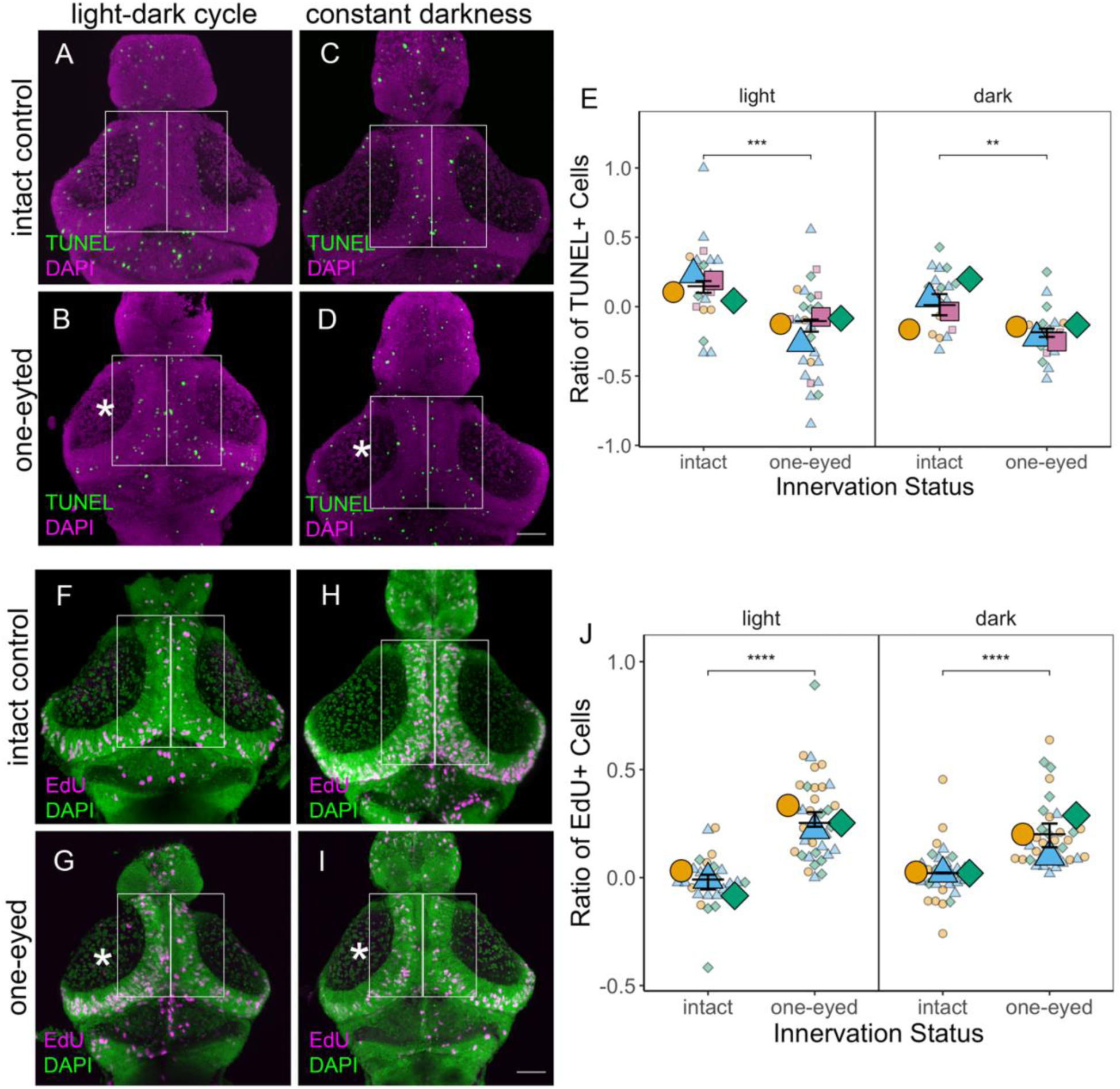
Cell survival and proliferation in the tectum is not significantly altered by restricting visual experience by dark rearing. **(A-D)** Maximum intensity projections of dorsal views of dissected brains showing TUNEL+ cells (green) and DAPI-stained nuclei (magenta) in the optic tecta of intact (A, C) and one-eyed (B, D) larvae raised in a typical light-dark cycle (A, B) or constant darkness (C, D) until 9 dpf. Innervated tectal lobes indicated with an asterisk. Bar is 50 µm. TUNEL+ cells inside boxed regions, excluding those in the tectal neuropil, w re counted and compared. **(E)** Superplot comparing the relative number of TUNEL+ cells in left (innervated post-extirpation) versus right (denervated post-extirpation) tectal lobes across four small batches of samples. Distinct colors and shapes indicate each batch; larger shapes represent median value of each batch; smaller shapes are individual data points (total n ≥ 18 for each condition). Abbreviations: ** p ≤ 0.01, *** p ≤ 0.001 (ANOVA followed by Tukey HSD for multiple comparisons). **(F-I)** Maximum intensity projections of dorsal views of dissected brains showing EdU+ cells (magenta) and DAPI-stained nuclei (green) in the optic tecta of intact (F, H) and one-eyed (G, I) larvae raised in a typical light-dark cycle (F, G) or constant darkness (H, I) until 9 dpf. Innervated tectal lobes indicated with an asterisk. Bar is 50 µm. EdU+ cells inside boxed regions, excluding those in the neuropil, were counted and compared **(J)** Superplots comparing the relative number of EdU+ cells in left (innervated post-extirpation) versus right (denervated post-extirpation) tectal lobes across three small batches. Distinct colors and shapes indicate each batch; larger shapes represent median value of each batch; smaller shapes are individual data points (total n ≥ 22 for each condition). Abbreviations: **** p ≤ 0.0001 (ANOVA followed by Tukey HSD for multiple comparisons)\

Comparing the ratios of TUNEL+ cells in one-eyed and intact larvae revealed that innervation status was the main contributor to the differences observed (ANOVA, f(1)=22.162, p<0.0001; ω^2^=0.57 [0.21, 1.00]). There was no statistically significant effect of environmental light condition (ANOVA, f(1)=3.078, p>0.11) and no interaction effect between innervation status and light condition (ANOVA, f(1)=0.553, p>0.47). Pairwise comparisons confirmed that, as in Fig 2, one-eyed fish reared in a typical light-dark cycle contained more TUNEL+ cells in the denervated side of the OT (Fig 4E; Tukey HSD, p<0.001). The same trend of more TUNEL+ cells on the denervated side of the OT was observed when examining one-eyed fish reared in near constant darkness (Fig 4E; Tukey HSD, p<0.024). Moreover, one-eyed larvae reared in either typical light-dark or near constant dark conditions did not exhibit statistically significant differences in the proportion of dying cells on the denervated versus innervated sides (Fig 4E; Tukey HSD, p>0.9). The difference (effect size) between the mean ratios of TUNEL+ cells in the OT of intact versus one-eyed larvae in the light condition (∂ = 0.279) was slightly larger than the difference between the mean ratios of intact versus one-eyed larvae reared in constant darkness (∂ = 0.204), but this difference fell just short of statistical significance (ω^2^ = −0.03 [0.00-1.00]).

Comparing the ratios of EdU+ cells in one-eyed and intact larvae showed that innervation status strongly correlated with proliferation (ANOVA, f(1)=40.594, p<0.001; ω^2^=0.77 [0.42, 1.00]) regardless of environmental light condition. There was no statistical difference between levels of innervation-dependent proliferation in light or dark-reared larvae (ANOVA, f(1)=0.190, p=0.67) and no interaction effect between innervation status and light condition (ANOVA, f(1)=2.537, p=0.14). Pairwise comparisons indicate that innervated tectal lobes contained more EdU+ cells, both in typical light-dark (Tukey HSD, p<0.001) and near constant dark (Tukey HSD, p<0.001) rearing conditions. As with cell death, dark rearing seemed to decrease the average difference between the amount of proliferation in both innervated and denervated tectal lobes, but not to a statistically significant degree (Fig 4J; ∂ = −0.289 for light-reared larvae and ∂ = −0.174 for dark-reared larvae; ω^2^ = 0.11 [0.00, 1.00]).

Dark rearing and innervation status contributed minimally to differences in brain and larval size. No statistically significant differences in OT volume were observed when comparing wild-type and one-eyed larvae (Fig S5 and Table S3). Very small differences in total body length of one-eyed larvae were observed in light but not dark rearing conditions (Fig S6 and Tables S3 and S4).

### Extent of *sox2+* expression correlates with innervation

To test whether a particular cell population was altered by innervation, we examined expression of *sox2*, a marker for neural stem and progenitor cells (reviewed in (Schmidt et al., 2013); see Fig 5A-D; F-I for representative images). Due to variability in staining intensity for *sox2* signal, we identified the volume of *sox2*-expressing cells relative to total DAPI-stained volume of the OT rather than counting discrete cells. Calculating this volumetric ratio allowed comparisons across all samples and revealed that both wild-type and innervated tecta contained a larger *sox2+* cell volume at 9 dpf (Fig 5E, J). Comparisons among wild-type and *lak* mutant tecta detected differences based on genotype (ANOVA, f(1)=16.824, p<0.001; ω^2^ = 0.25 [0.08, 1.00]) with pairwise comparisons identifying a statistically significant difference between *lak* mutant and wild-type OT only at 9 dpf (Tukey HSD, p<0.001). Comparisons among intact and one-eyed larvae provide support for innervation (ANOVA, f(1)=43.82, p<0.001) and age (ANOVA, f(1)=25.02, p<0.001) both contributing to the extent of *sox2* expression. Pairwise comparisons detected statistically significant differences at 9 dpf, with the innervated sides of the tecta containing nearly twice the relative *sox2* volume as the denervated sides (Fig 5H-J; Tukey HSD, p<0.001). Together these results indicate that optic nerve innervation may promote maintenance and/or proliferation of *sox2*+ neural stem cells in the optic tectum.

**Figure 5:**
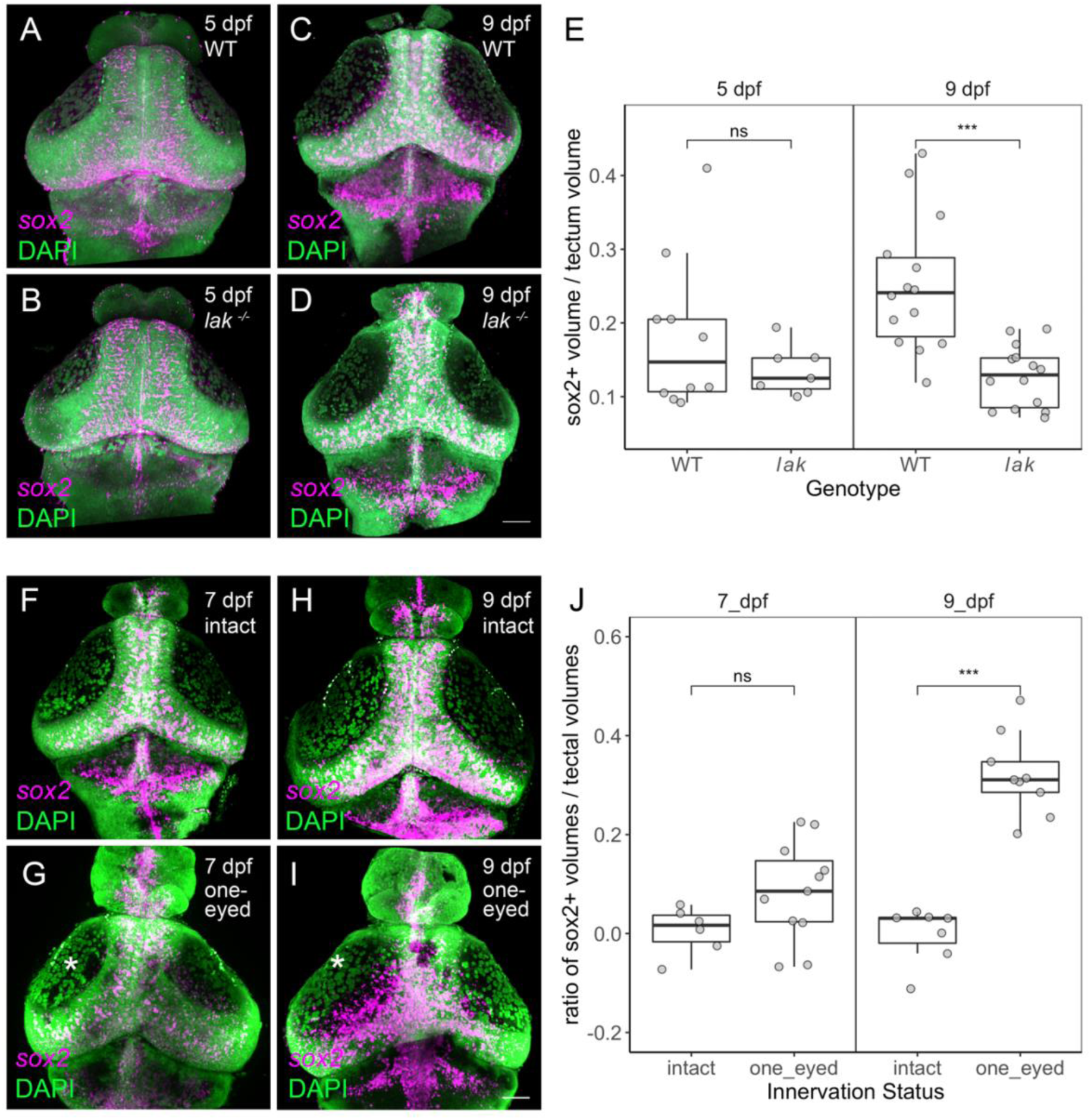
*sox2+* stem cells are depleted in tecta lacking optic nerve innervation. **(A-D)** Maximum intensity projections of dorsal views of dissected brains from wild-type (A, C) and *lak^-/-^* (B, D) larvae showing *sox2* expression (magenta) by fluorescent in situ hybridization with nuclear counterstain (DAPI, green). Bar is 50 µm. Ages and genotypes indicated in the upper right corner of all panels. **(E)** Box plots overlaid with individual data points depict the volumetric ratios of *sox2*-expressing regions relative and DAPI-stained volumes in dorsal midbrain at indicated ages. n ≥ 7 for each condition. Abbreviations: ns, not significant; *** p ≤ 0.001 (Wilcoxon ranked sums test). **(F-I)** Maximum intensity projections of dorsal views of dissected larval brains from intact (F, H) and one-eyed (G, I) larvae showing *sox2* expression (magenta) by in situ hybridization with nuclear counterstain (DAPI, green). Bar is 50 µm. Innervated tectal lobes indicated with an asterisk. Ages and surgery status indicated in the upper right corner of all panels. **(J)** Box plots overlaid with individual data points depict volumetric ratios of *sox2*-expressing regions relative to the nuclear volume of the corresponding (left (innervated after extirpation) versus right (denervated after extirpation) tectal lobes at indicated ages. n ≥ 6 for each condition. Abbreviations: ns, not significant; *** p ≤ 0.001 (Wilcoxon ranked sums test).

### Wnt/β-catenin pathway is upregulated in the OT with retinal innervation

To address some of the molecular cues that enable the optic nerve to signal proliferation and survival of stem and progenitor cells in the OT, we examined expression of genes encoding components of different signaling pathways known to be involved in proliferation in the visual system. The Sonic hedgehog pathway (Shh) has previously been shown to promote cell proliferation and *sox2* transcription in the adult zebrafish OT (Shitasako et al., 2017). We found, however, that components of the Hedgehog signaling pathway did not appear to exhibit any noticeable differences in expression between innervated and denervated tectal lobes in embryonic and larval zebrafish (Fig S7).

The Wnt/β-catenin pathway is required for a multitude of events throughout all stages of vertebrate nervous system development including patterning and morphogenesis, proliferation and differentiation, synapse formation and plasticity, and maintenance of pluripotency (Bielen and Houart, 2014; Faraji et al., 2025; Mulligan and Cheyette, 2012). In adult zebrafish, Wnt/β-catenin pathway activity in the OT promotes regeneration after acute injury (Shimizu et al., 2018; Shitasako et al., 2017). In larval zebrafish brains, we found that expression of *axin2,* a Wnt/β-catenin pathway target, was innervation dependent (Fig 6). To test whether the Wnt/β-catenin pathway was differentially activated in innervated tectal lobes, we performed in situ hybridization for *axin2* and *lef1* on brains isolated from intact, two-eyed larvae and one-eyed larvae generated by unilateral eye extirpation at 5 dpf. Comparisons of innervated and denervated patterns showed that innervation status positively correlated with the extent of *axin2* (p<0.001) and *lef1* (p=0.015) expression (Fig 6E). In particular, innervated OT lobes contained nearly twice as many *axin2+* cells as denervated OT lobes (Fig 6B and 6B’). Reverse-transcription quantitative PCR (qPCR) using cDNA from samples collected from innervated and denervated tectal lobes corroborated significantly higher levels of *axin2* when OT lobes were innervated (Fig 6F; *axin2*, p=0.01), but the change in levels of *lef1* between innervated and denervated OT lobes did not reach statistical significance (Fig 6F; lef1, p=0.14). Genes including *ccnD1* (p=0.31) and calibrator gene *ef1a* were expressed at similar levels, irrespective of optic nerve innervation (Fig 6F).

**Figure 6:**
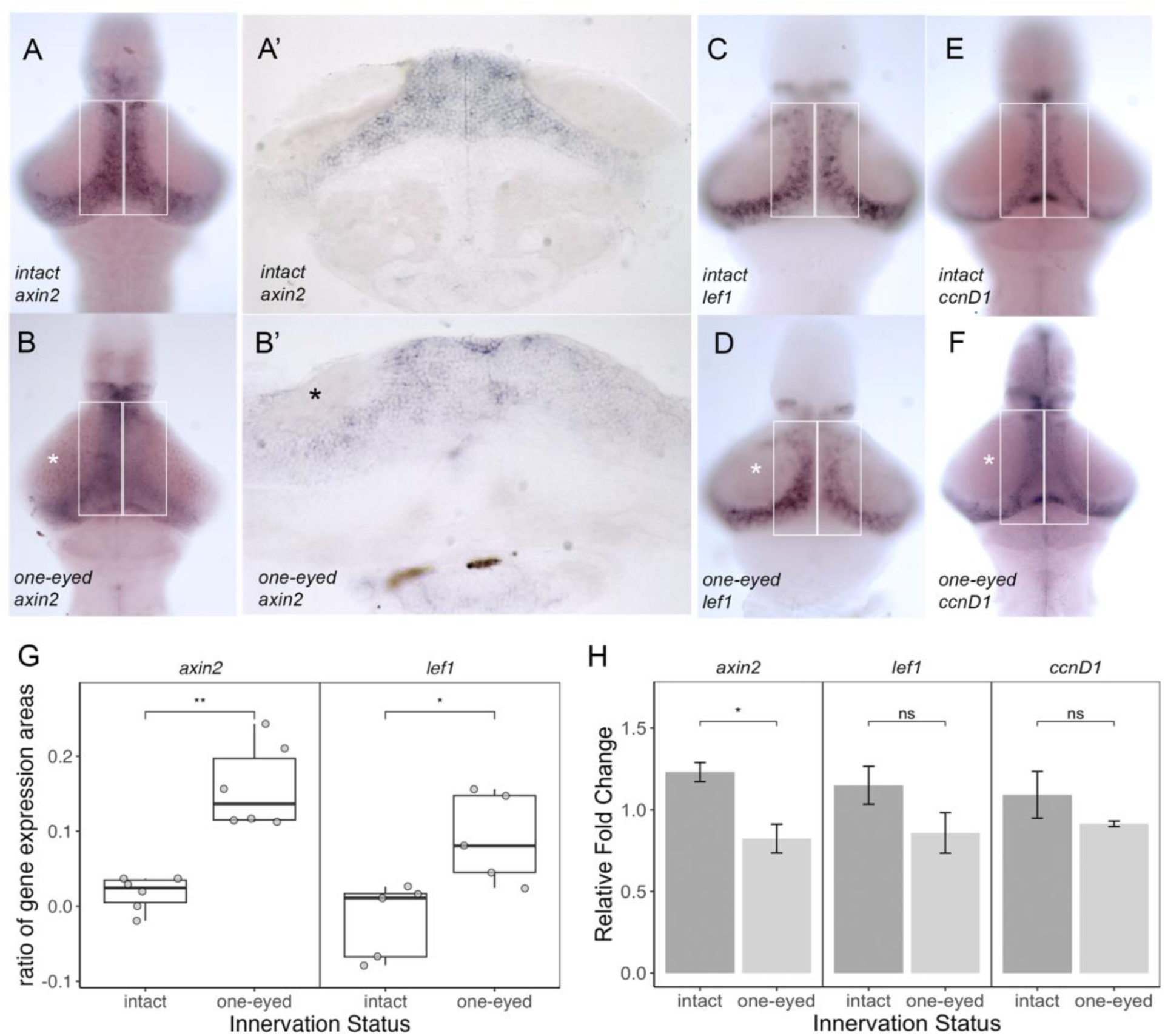
Wnt/B-catenin pathway target genes show decreased tectal expression when innervation is removed. **(A-F)** Expression of Wnt/B-catenin target genes, *axin2* (A-B) and *lef1* (C-D), and control gene *ccnD1* (E-F), as revealed by chromogenic in situ hybridization on whole mount brains from intact (A, C, E) or one-eyed (B, D, F) 9 dpf larvae. Boxed regions on dorsal views of whole mount brains indicate the region in which gene expression area was quantified. Thin frontal sections of *axin2*-stained brains from intact (A’) and one-eyed (B’) midbrains. Innervated tectal lobes indicated with asterisk. Target gene and surgery status indicated in lower left corner of panels. **(G)** Box plots overlaid with individual data points showing ratios of the gene expression areas (left or innervated – right or denervated) / total area for *axin2* or *lef1*), n>5 for each condition and gene. Abbreviations: ns, not significant, * p ≤ 0.05, ** p ≤ 0.01 (Wilcoxon ranked sums test). **(H)** Bar plot showing mean relative fold change in expression of *axin2*, *lef1*, and *ccnD1* by qRT-PCR as normalized to *ef1a*. Bars = standard error. n = 4 biological replicates of ∼20 dissected larval midbrains per condition per replicate. Abbreviations: ns, not significant; * p ≤ 0.05 (Welch’s t-test).

Similar to denervated OT lobes, *lak* mutant OT showed decreased *axin2* expression as revealed by *in situ* hybridization (Fig 7A-B, E; p<0.001) and qPCR (Fig 7M; p<0.002). Unlike the denervated OT lobes, *lak* mutant OT did not show a statistically significant decrease in *lef1* expression region by *in situ* hybridization (Fig 7C-D, E; p<0.077). Given Wnt/β-catenin pathway activity is linked to *sox2* levels in cultured stem cells (Blassberg et al., 2022; Mukherjee et al., 2022) and the decreased number of *sox2+* cells in denervated and *lak* mutant OT detected by *in situ* hybridization (Fig 5), we examined *sox2* expression by qPCR. Buttressing our *in situ* hybridization data and found decreased levels of *sox2* mRNA in *lak* mutants (Fig 7M; p<0.04).

**Figure 7:**
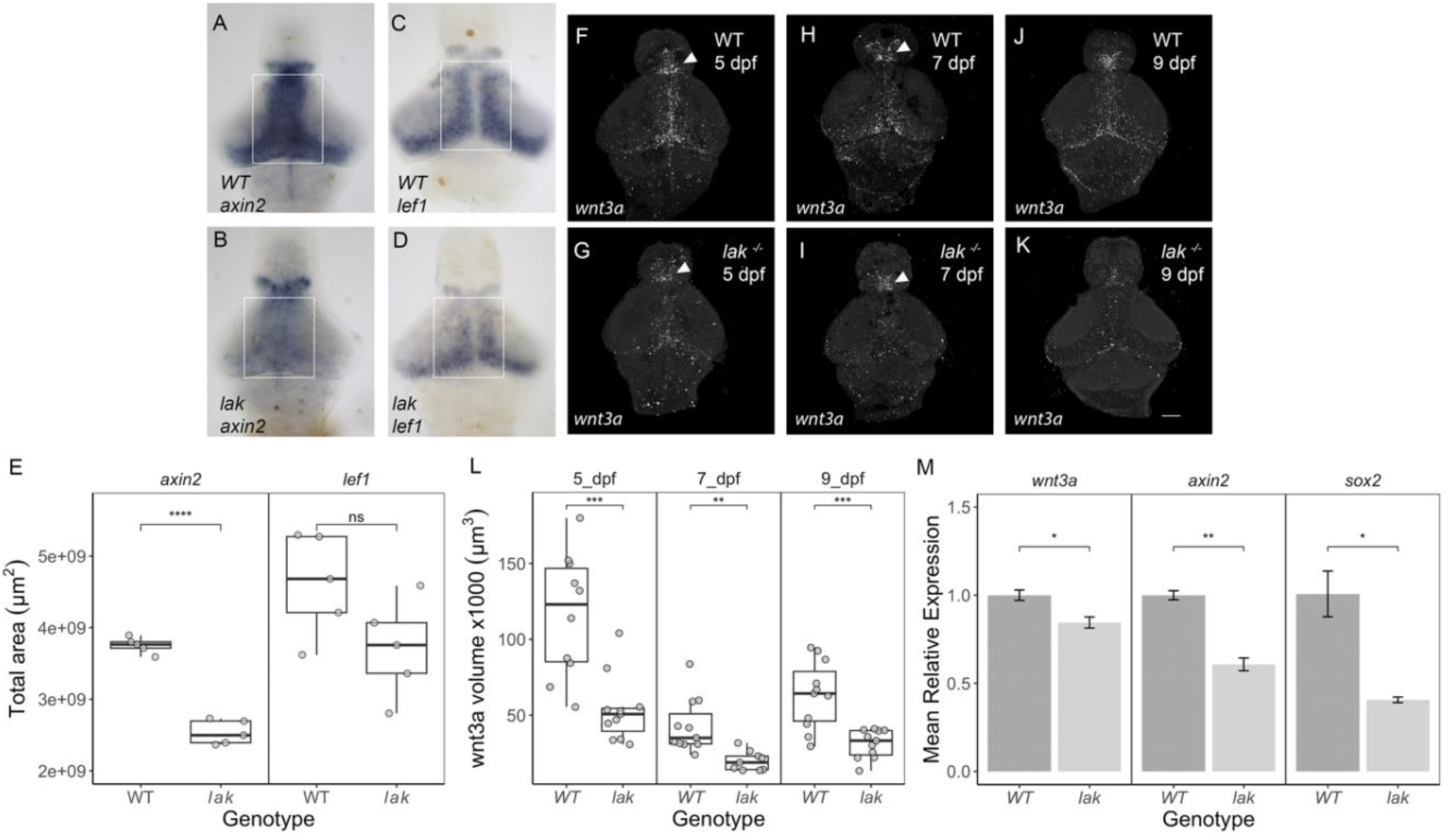
Wnt3a and Wnt/β-catenin target gene expression is reduced in the OT when optic nerve innervation is absent. **(A-D)** Dorsal views of whole mount dissected brains showing *axin2* (A, B) or *lef1* (C, D) expression by chromogenic in situ hybridization of wild-type (WT; A, C) and *lak^-/-^* (B, D) larvae. Boxes indicate the region in which gene expression area was quantified. Genotype and target gene indicated in the lower left corner of all panels. **(E)** Box plots overlaid with individual data points show total area positive for expression of indicated gene. n = 5 for each condition. Abbreviations: ns, not significant; *** p ≤ 0.001 (Wilcoxon ranked sums test). **(F-K)** Maximum intensity projections of dorsal views of dissected brains from wild-type (F, H, J) and *lak^-/-^* (G, I, K) larvae highlighting *wnt3a* transcripts (white puncta) detected with fluorescent in situ hybridization. Bar is 50 µm. Arrow heads indicate *wnt3a*-expressing regions in the forebrain. Ages and genotype indicated in upper right corner of all panels. **(L)** Box plot overlaid with individual data points showing *wnt3a* expression volumes for ages and genotypes indicated. n ≥ 9 for each condition. Abbreviations: ** p ≤ 0.01, *** p ≤ 0.001 (Wilcoxon ranked sums test). **(M)** Bar plot showing mean relative expression of *wnt3a*, *axin2*, and *sox2* by qRT-PCR as normalized to *actb1*. Bars = standard error. n = 3 biological replicates of ∼20 dissected larval brains per condition per replicate. Abbreviations: * p < 0.05, ** p ≤ 0.01 (Welch’s t-test).

Although a number of *wnt* genes are expressed in the dorsal midbrain during early development (Farnsworth et al., 2020; Sur et al., 2023), *wnt3a* has been shown to be robustly expressed in the dorsal midbrain and optic tectum at later stages of development (3 dpf; (Duncan et al., 2015) and 6 dpf; (Sherman et al., 2023)). We probed wild-type and *lak* mutant brains for *wnt3a* expression (Fig 7F-K) with fluorescent in situ hybridization and found that wild-type tecta contained greater volumes of *wnt3a+* cells at 5, 7, and 9 dpf (Fig 7L). Comparisons across ages and genotypes indicated that both variables contribute to differences in *wnt3a*

mRNA expression (ANOVA ages, f(2)=30.529, p<0.001; ANOVA genotype, f(1)=42.339, p<0.001) with age and genotype exhibiting a small but significant interaction effect (ANOVA, f(2)=4.388, p=0.017; ω^2^=0.04 [0.00-1.00]). Pairwise comparisons showed that *lak* mutants contained smaller *wnt3a*-expressing regions, particularly at 5 dpf (Tukey HSD, p<0.001) and 9 dpf (Tukey HSD, p=0.018). Additional pairwise comparisons showed that at 5 dpf, wild-type tecta contained significantly larger *wnt3a*-expressing regions than at 7 dpf (Tukey HSD, p<0.001) or 9 dpf (Tukey HSD, p<0.001). The presence of *wnt3a* transcripts in the forebrain appears generally unchanged in *lak* mutants, especially at 5 and 7 dpf (arrow heads in Fig 7F-I). Consistent with the *in situ* hybridization data, qPCR of cDNA from 9 dpf wild-type and *lak* mutant brains also revealed a significant decrease in *wnt3a* expression (Fig 7M; p=0.023).

### Wnt/β-catenin pathway activation restores levels of tectal cell proliferation in absence of retinal innervation

If Wnt pathway activity promotes proliferation in an innervation-dependent manner, then activating the Wnt/β-catenin pathway in *lak* mutants could restore proliferation close to wild-type levels. To test this hypothesis, clutches of fish from a *lak*^+/-^ incross were treated with 1 µM BIO, a Wnt/B-cat pathway activator, starting at 7 dpf and then incubated in BIO and EdU at 8.5 dpf so that the larvae were exposed to BIO for ∼48 h and EdU for ∼16 h (Fig 8A-D). Analysis of EdU incorporation indicated that although genotype was the main contributor to observed differences in proliferation (ANOVA, f(1)=14.145, p<0.001), genotype and BIO treatment exhibited a significant interaction effect (ANOVA, f(1)=4.597, p=0.036). Pairwise comparisons confirmed that, as in Fig 3, *lak* mutant tecta contained fewer proliferative cells (Tukey HSD, p<0.001).

**Figure 8:**
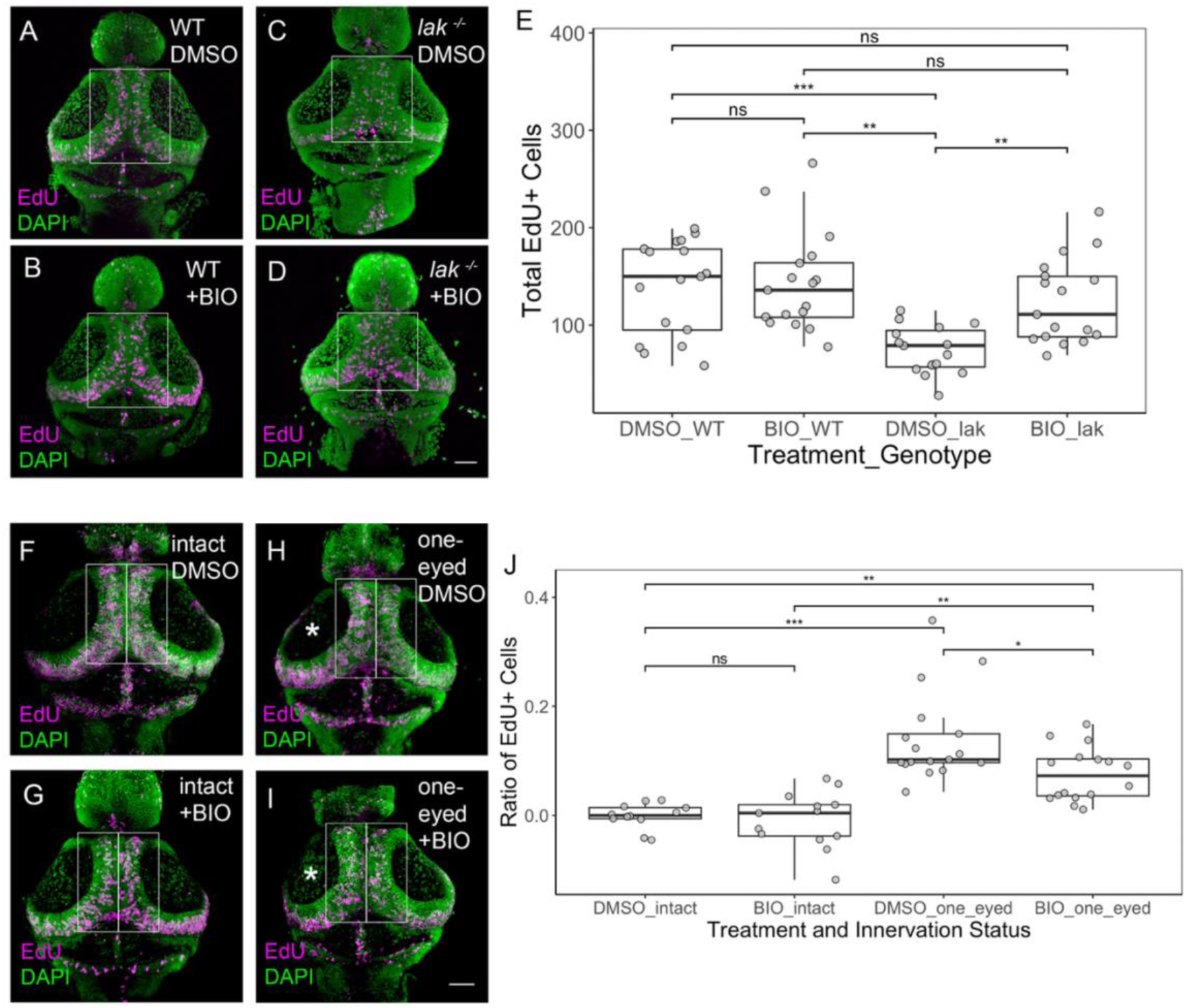
The Wnt/B-catenin pathway agonist, BIO, restores tectal cell proliferation in mutants lacking innervation. **(A-D)** Maximum intensity projections of dorsal views of dissected brains from wild-type (A, B) and *lak^-/-^* (C, D) larvae treated with 1µM 6-bromoindirubin-3-oxime (BIO; B, D) or an equivalent volume of vehicle control (DMSO; A, C) showing EdU+ cells (magenta) with nuclear counterstain (DAPI, green). Bar is 50 µm. Boxes indicate the region in which EdU+ cells were counted. Genotype and treatment indicated in upper right corner of all panels. **(E)** Box plot overlaid with individual data points indicating number of EdU+ cells as a function of genotype and treatment; n ≥ 15 for each condition. Abbreviations: ns, not significant; ** p ≤ 0.01; *** p ≤ 0.001 (ANOVA followed by Tukey HSD for multiple comparisons). **(F-I)** Maximum intensity projections of dorsal views of dissected brains from intact (F, G) and one-eyed (H, I) larvae treated with 1µM BIO (G, I) or an equivalent volume of vehicle control (DMSO; F, H) showing EdU+ cells (magenta) with nuclear counterstain (DAPI, green). Bar is 50 µm. Boxes indicate the region in which EdU+ cells were counted. Asterisks mark innervated tectal lobes. Surgery status and treatment indicated in upper right corner of all panels. **(J)** Box plot overlaid with individual data points indicating the ratio of EdU+ cells on the left (innervated post-surgery) relative to right (denervated post-surgery) side of the OT in each condition; n ≥ 12 for each condition. Abbreviations: ns, not significant; * p ≤ 0.05; ** p ≤ 0.01; *** p ≤ 0.001 (ANOVA followed by Tukey HSD for multiple comparisons).

**Figure 9:**
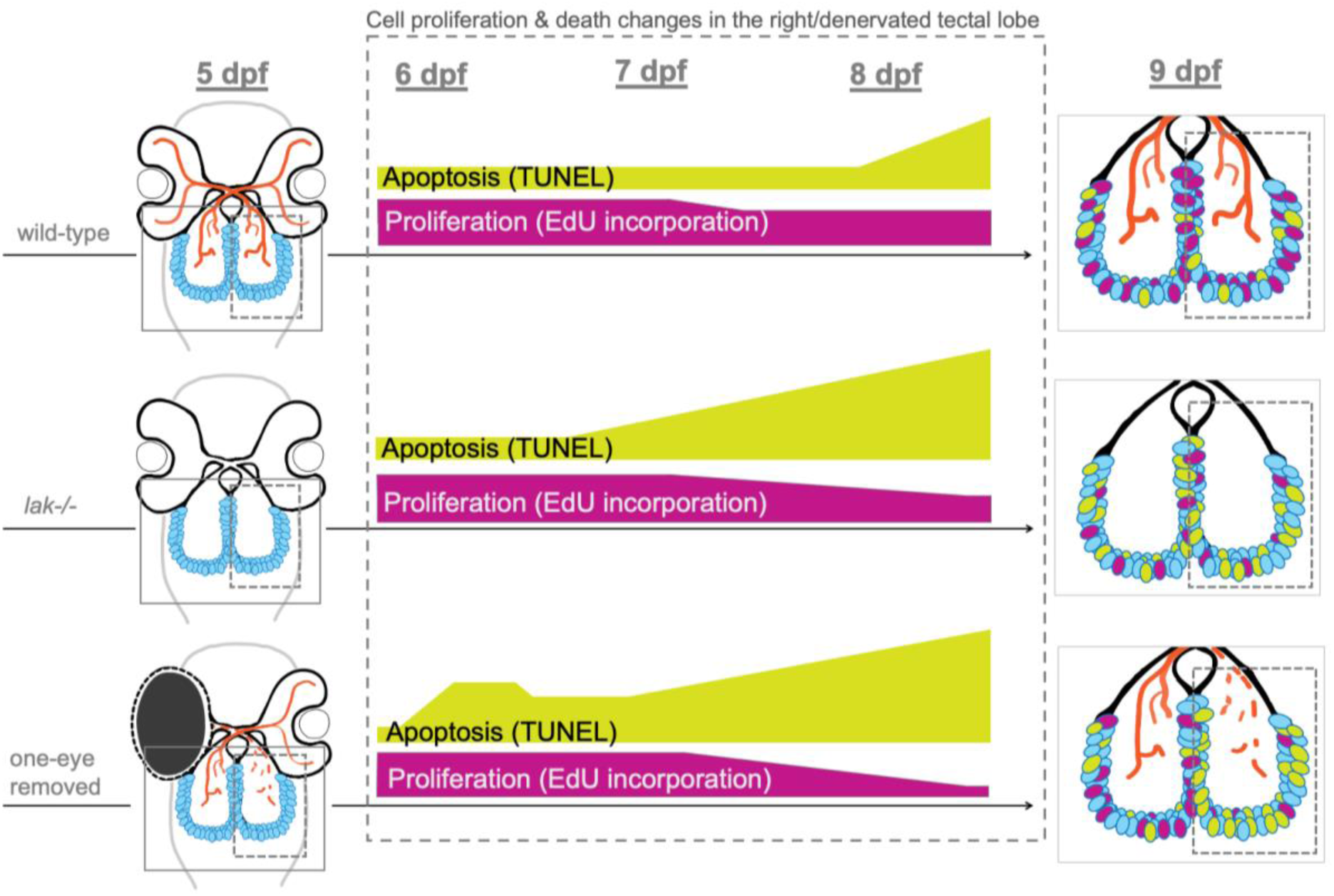
Summary of results showing correlation between Wnt/B-catenin pathway activity, optic nerve innervation, cell proliferation, and death in the optic tectum. Diagrammatic summary of changes in cell death (TUNEL, yellow) and proliferation (EdU incorporation, magenta) for boxed regions of optic tectum in wild-type, *lakritz* (*atoh7-/-*), and denervated (one-eye removed) larvae from 6 to 9 dpf.

However, when *lak* mutant larvae were incubated with BIO, the average number of EdU+ cells in the mutant OT (125 ± 44) was indistinguishable to DMSO-treated wild-type (Tukey HSD, p=0.78). In addition, the average number of EdU+ cells in control wild-type larvae (DMSO-treated) was 139 ± 48, which was not statistically significantly different from the average number of EdU+ cells in BIO-treated (143 ± 51) wild-type larvae (Tukey HSD, p=0.99).

To determine if exogenous activation of the Wnt/β-catenin pathway could also rescue cell proliferation in dennervated OT, we treated intact larvae and those that had one eye removed at 5 dpf with BIO and EdU as above (Fig 8F-J). The numbers of EdU+ cells within the midline region of each side of the TPZ (e.g., boxes in Fig 8F-I) were counted and compared by calculating the difference between the number of EdU+ cells in the innervated (or left) side and denervated (or right) side and then dividing by the total number of EdU+ cells. As in Fig 3, this difference ratio is close to zero when the number of proliferating cells is similar in both left and right OT lobes (e.g., intact larvae; Fig 8F-G, J). When the denervated OT sides have fewer EdU+ cells, the difference ratio is greater than zero (e.g., one-eyed larvae; Fig 8H-I, J). Analysis of EdU incorporation indicated that although innervation status was the main contributor to observed differences in proliferation (ANOVA, f(1)=52.945, p<0.001), innervation status and BIO treatment exhibited a significant interaction effect (ANOVA, f(1)=8.314, p<0.006). Pairwise comparisons confirmed that, as in Fig 3, denervated OT contained fewer proliferative cells as compared to innervated intact OT (Tukey HSD, p<0.001). Supporting links among innervation,

Wnt/B-pathway activity, and proliferation, we found that the ratios of EdU+ cells in the OT of one-eyed BIO-treated larvae were significantly lower (closer to zero) than the ratios of EdU+ cells in the OT of one-eyed DMSO-treated larvae (Tukey HSD, p=0.012). BIO treatment did not, however, completely equalize the number of EdU+ cells in denervated and innervated OT. The difference ratio of EdU+ cells in these two populations was still significantly different (Tukey HSD, p<0.007). Although these data support a role for innervation-dependent Wnt/β-catenin regulated growth in the optic tectum, they also indicate that injury-induced loss of retinal innervation may elicit different molecular signals and cell behaviors as compared to the absence of retinal innervation during all stages of development.

## Discussion

While all vertebrates exhibit some life-long remodeling and growth in their nervous systems, zebrafish support extensive life-long growth with specialized stem and progenitor cells arrayed throughout the nervous system (Becker and Becker, 2022; Foley et al., 2024). In this study, we set out to investigate how innervation from the zebrafish eye could regulate growth and development of its recipient tissue in the brain. With a combination of surgical and genetic perturbations to remove retinal ganglion cell input into the optic tectum, our data provide evidence that progenitor cell survival and proliferation requires innervation-dependent Wnt/β-catenin pathway activity in the optic tectum only after it and its connections with other regions are established. Our work lays the foundation for future experiments that will unravel connections between sensory system input and Wnt/β-catenin pathway activity in the growing and developing brain.

When we tested whether innervation was required for the survival and proliferation of stem and progenitor cells in the optic tectum, we found that cells in the visual processing regions of the zebrafish brain initially follow an intrinsic program for their development.

Specifically, we find no change in the number of proliferating or dying cells in tectal lobes of wild-type or *lak* mutant zebrafish until ∼7 dpf. A recent single nuclear RNA Sequencing (snRNA-Seq) analysis of samples from wild-type and *lak* mutant brains at 6 dpf used pseudotime and RNA-velocity analyses to estimate the proportions of cells in distinct cell cycle phases, suggesting that progenitor cells in *lak* OT cycle and differentiate more slowly than cells in wild-type OT (Sherman et al., 2023). While our EdU to analyses could also be also consistent with a longer cell cycle for progenitor cells in the *lak* tectum, lengthening cell cycle time is not the only explanation for fewer proliferating cells. Both elevated levels of programmed cell death in as well as a possible decreased probability of entering the cell cycle in *lak* mutant tecta, could also contribute to cell survival and proliferation in the OT downstream of optic nerve innervation.

Our results also highlight several underappreciated key features of optic tectum development, including a time window for programmed cell death in zebrafish brain development. In contrast to mammalian brains, zebrafish brains exhibit limited developmental apoptosis, especially when the progeny of neural progenitors remain near their site of generation (Zupanc et al., 2009). To date, most studies have examined larvae at 5 and 6 dpf or 14+ days later, with little evidence that apoptosis plays a significant role in sculpting visual system development. By extending the timeframe of analyses, we identify 7-9 dpf as a critical window during which optic nerve innervation contributes to cell survival and proliferation. As this stage occurs after RGC-OT connections are well established, the local environment of the OT may change such that after 6 dpf apoptosis may be used to refine synaptic connections as has been documented in a variety of other organisms, including mammals and invertebrates (Ryu et al., 2016). Many open questions about how programmed cell death contributes to the neuronal circuit remodeling required for lifelong growth and development.

It is also unclear whether and how connections other than those from the retina influence optic tectum growth and development. For instance, RGC axons from the optic nerve converge on two brain regions -- nuclei within the thalamus (forebrain) and the OT/superior colliculus in the dorsal midbrain. Although different vertebrates send different proportions of RGC axons to these distinct locations, retino-thalamic and retio-colliculo-thalamic projections are conserved among all vertebrates examined to date (Isa et al., 2021). Interestingly, a study examining how loss of RGC innervation of the dorsal lateral geniculate nucleus (dLGN) in mammals provides evidence that cell survival in the dLGN is more dependent on efferent connections to the visual cortex rather than afferent input from the retina (Zacharaki et al., 2010). It is currently unknown how efferent connections to the circuits that evoke visually-guided behaviors in fish contribute to cell survival in the OT and/or OT growth and development.

Our results begin to provide a molecular connection between optic nerve innervation and proliferation, revealing that optic nerve innervation is connected with robust expression of *wnt3a* and activation of the Wnt/β-catenin pathway. Consistent with published sc-RNA sequencing experiments comparing wild-type and *lakritz* mutants (Covello et al., 2020; Sherman et al., 2023), we find that OT lacking innervation exhibit lower levels of *wnt3a* gene expression and concomitant decreases in Wnt/β-catenin pathway targets. Wnt pathway activity regulates neural stem cell behaviors in a variety of contexts and organisms with implications for brain development and homeostasis (e.g., Donval et al., 2024; Kalamakis et al., 2019; Sebo et al., 2025). Recently, *sox2* has been shown to direct B-catenin to enhancers of neural genes in a *lef/tcf*-independent manner (Mukherjee et al., 2022). Consistent with this published work, our data show reduced levels of *sox2* and *wnt3a* expression in *lak* mutants, raising the possibility that Wnt/β-catenin activity supports proliferation of *sox2+* progenitors in OT progenitors in response to optic nerve innervation.

Wnt/β-catenin pathway activity is linked to stem and progenitor cell proliferation in many contexts including the nervous system (Nusse and Clevers, 2017; Steinhart and Angers, 2018), and our data now suggest that innervation-dependent Wnt/β-catenin pathway activity could be one way to coordinate life-long growth of the retina and its target tissue in the brain. Not only do our results show decreased *wnt3a* expression in OT without innervation, they also illustrate that proliferation in the OT can be rescued by precisely-timed pharmacological activation of the Wnt/β-catenin pathway with the GSK3B-inhibitor, BIO. In addition to our results showing expression of *wnt3a* in stem and progenitor cells in the OT, a thorough search of publicly available scRNASeq datasets (e.g., (Farnsworth et al., 2020; Sur et al., 2023)) identifies expression of multiple *wnt* genes -- *wnt11r*, *wnt7ba*, *wnt8b*, *wnt4a*, *wnt10b* -- in proliferating midbrain progenitors during multiple stages of development. Determining whether all of these genes are co-expressed in the same cells and/or responsive to optic nerve innervation in tandem with elucidating the pathways and molecules downstream of Wnt/β-catenin in the OT, will provide new insights into innervation-dependent growth in vertebrate sensory systems.

It remains unclear how exactly RGCs activate OT progenitor cell proliferation through the Wnt/β-catenin pathway. One study has linked release of glutamate from RGCs in the optic tectum to cell proliferation showing that when zebrafish larvae were injected with APV, an NMDA glutamate receptor antagonist in their OT, cell proliferation appeared to decrease (Hall and Tropepe, 2018). In contrast to our studies showing negligible differences in cell proliferation and cell survival when fish were reared under no/low light conditions, Hall and Tropepe (2018) showed that wild-type fish reared in low-light exhibited decreased EdU incorporation in the OT. It is possible that differences in labeling approaches and/or image analysis could explain the discrepancies between the two studies. It is also possible that we missed small regional differences in our analyses of whole mount brains that their study detected by examining cryosections of subsections of the OT. Importantly, connections between neuronal electrochemical activity and neurogenesis have been documented in a variety of vertebrates, with studies implicating the release of brain derived neurotrophic factor (BDNF) with adult neurogenesis (Gage, 2025; Sato et al., 2016). One study of exercise-induced neurogenesis in the mammalian hippocampus provides some evidence that BDNF may collaborate with the Wnt/β-catenin pathway to enhance neurogenesis after injury (Cheng et al., 2020). In our study, we show a slight enhancement of proliferation when denervated tectal lobes are treated with BIO to activate the Wnt/β-catenin pathway in a different type of injury context.

Situating our study in the current literature illustrates that there are many opportunities to explore the cellular and molecular mechanisms that guide life-long neurogenesis, from both developmental and injury contexts. Future experiments exploring the roles of RGC and tectal cell electrochemical activity could provide more and deeper cell and molecular insights into how continuously integrating new input from the retina can regulate brain growth and development.

## Methods

### Fish Strains, Care, and Manipulations

Adult zebrafish (*Danio rerio*) were maintained on a 14-hour light (from 200 to 246 lx) /10-hour dark cycle at 26.5°C - 28°C (Westerfield, 2007). Wild-type (WT) embryos were obtained from crossing AB and Tübingen strains (Zebrafish International Resource Center).

Heterozygous transgenic and mutant carrier lines (*Tg[atoh7:RFP] (Poggi et al., 2005)* and *lakritz^th241^* (Kay et al., 2001) were crossed to generate embryos and larvae; progeny from *lakritz^+/-^ ;Tg[atoh7:RFP]* were sorted into mutants and siblings at 5 days post fertilization (dpf) based on the presence of a fluorescent optic tract or genotyped as described (Kay et al., 2001).

Embryos were obtained by natural spawning and then transferred into embryo medium (E3;((Nusslein-Volhard and Dahm, 2002)). E3 was changed daily and after 5 dpf, larvae were fed rotifers every 24 hours. Larvae were humanely culled with a lethal dose of buffered tricaine at 5 dpf, 6 dpf, 7 dpf, or 9 dpf. E3 was supplemented with 200 µM PTU to diminish melanin synthesis (Westerfield, 2007).

For experiments that required dark rearing, embryos were separated into light and dark treatment groups before 8 hours post-fertilization (hpf). Light-reared embryos were raised in the fish room under the same conditions as adults. Dark-reared embryos were raised in a light-tight box constructed from sheet metal and fitted around one row of a single-sided Aquaneering zebrafish rack, keeping out all detectable light. During the enucleation procedure, dark-reared embryos, including the non-enucleated controls, were exposed to low ambient light (∼142 lx) for approximately 2 hours. On the days following enucleation, dark-reared embryos were exposed to minimal low light (8-12 lx) for about 5 minutes per day during water changes, and an additional 5-10 minutes of ambient light when EdU was added to their water. This level of light is in stark contrast to the duration and intensity of light (200-246 lx for 14 h per day) that control (light-reared) embryos experienced.

For experiments comparing innervated and non-innervated tectal lobes, 5 dpf larvae were anesthetized in E3 with 0.015% buffered tricaine until they were non-responsive to touch. They were then embedded in small drops of 1.5% low melting point (LMP) agarose made in Marc’s Modified Ringer’s Solution (MMR) with their left side facing upward. The left eye was loosened and removed with an electrolytically sharpened tungsten needle (Hagen et al., 2022; Turner et al., 2014). The larvae were then freed from the agarose and incubated overnight in 1X MMR supplemented with 1% penicillin/streptomycin (P/S, Sigma). Larvae were transferred to E3 and fed rotifers 1 day after surgeries were performed.

To activate the Wnt/β-catenin pathway, at 7 dpf larvae were treated with either 1 µM 6-bromoindirubin-3-oxime (BIO; Sigma) diluted into E3 from a 1mM stock made in DMSO or with an equivalent volume of DMSO. The same treatment was reapplied at 8 dpf.

All experiments were performed in accordance with guidelines and protocols approved by the Institutional Animal Care and Use Committees of Reed College and UCL.

### Brain dissection and histochemistry

After anesthetization with a lethal dose of tricaine (Matthews and Varga, 2012), larvae were fixed in 4% paraformaldehyde (PFA) supplemented with 4% sucrose overnight at 4°C, and washed into phosphate buffered saline (PBS) the next day. Fixed larvae were suspended in

PBS droplets and secured laterally to a Sylgard (Sylgard 184, Dow) plate using tungsten pins through the notochord. A sharpened tungsten needle and needle-sharp forceps were used to expose the brain by removing, in order, the eyes, ear, jaw, gut, dorsal cranial skin, and palate. Brains were placed in fresh PBS at 4°C following dissection, until they were gradually dehydrated into 100% methanol (MeOH). Brains were stored at −20°C in 100% MeOH until they were subjected to EdU detection, TUNEL, *in situ* hybridization, or immunohistochemistry. Many of the brains that retained melanin pigment after dissection were bleached by incubating them in a solution of 1% KOH (diluted from a freshly made 10% stock solution) and 1.2% H_2_O_2_ (diluted from 30% concentrated stock) in PBS for 5-10 minutes at room temperature. Embryos were rinsed 3-5 times in PBS + 0.1% TritonX-100 (PBST) to stop the reaction.

Proliferating cells were detected by first swimming larvae in 5-ethynyl-2’-deoxyuridine (EdU, Invitrogen). A 6.7 mM EdU stock solution was made in a 2:1 mix of E3:DMSO and stored at −20°C for weeks at a time. Larvae were transferred to 35 mm petri dishes containing 0.4 mM EdU in E3 and incubated for 16-18 hours in their assigned light/dark condition or for briefer pulses as described. EdU treatment was replaced with E3 until fish were fixed (typically 1-3 hours later but for pulse-chase experiments up to 4 days later). Following brain dissection, EdU-positive cells were detected with the Click-It EdU Alexa Fluor 647 or 555 Imaging Kit (Invitrogen) according to the manufacturer’s instructions.

Apoptotic cells were detected using terminal deoxynucleotidyl transferase (TdT) dUTP nick end labeling (TUNEL; ApoTag reagents, Chemicon/Millipore) as previously described (Cerveny et al., 2010), except that dissected brains were treated with 40 µg/ml Proteinase K for 30 minutes at 37°C for permeabilization. TUNEL-positive cells were detected with either anti-Digoxigenin-AP Fab fragments (Roche, 1:3000) or anti-Digoxigenin-Fluorescein (Roche, 1:1000).

Whole-mount immunohistochemistry was performed essentially as previously described (Cerveny et al., 2010; Hagen et al., 2022). Larval brains were incubated with antibodies diluted in IB Buffer (PBS supplemented with 1% TritonX-100, 10% NGS, and 1% DMSO) for 18-20 hours at 4°C. Larvae were washed into PBS and stored at 4°C prior to imaging. Primary antibodies were chicken anti-GFP (1:500, Abcam) and rabbit anti-RFP (1:500, MBL); secondary antibodies were anti-chicken Alexa Fluor-488 and anti-rabbit Alexa Fluor-567 (both Molecular Probes, 1:500). For nuclear staining, larvae were incubated in DAPI (1 µg/ml) at 4°C overnight prior to imaging.

### Gene expression analysis

For *in situ* hybridization, assays were performed essentially as previously described (Xu et al., 1994). Digoxygenin-labled anti-sense RNA probes were synthesized as previously described for *axin2* (Weidinger et al., 2005)*, lef1* (Lee et al., 2006), and *wnt3a* (Duncan et al., 2015). A *sox2* probe was synthesized with T3 polymerase and template that was amplified from cDNA by the polymerase chain reaction using primers listed in Table S1. Gene expression patterns were detected with anti-Digoxygenin AP Fab fragments (Roche at 1:1000) and either NBT/BCIP (Roche), Fast Red (Sigma), or Fast Blue (Sigma) (Lauter et al., 2011; Moens, 2008).

For quantitative real time polymerase chain reaction (qRT-PCR) experiments, two approaches for isolating RNA were used. For *lak* mutant and wild-type, total RNA was extracted from sets of dissected brains from 9 dpf larvae grouped by genotype and fixed in 4% PFA in PBS with 4% sucrose. After fixation, brains were exposed through dissection (Hagen et al., 2022; Turner et al., 2014). For *lak* mutant and wild-type sibling larvae, whole brains were stored in Trizol-LS at −80°C after dissection. RNA was isolated from 3 sets of 20 brains for each condition using RNeasy columns (Qiagen) after chloroform extraction and its quality measured by OD260/OD280 nm absorption ratio > 2.0 and an OD260/230 nm absorption ratio between 1.98 - 2.12. cDNA was synthesized using iScript reverse transcriptase kit (BioRad).

For one-eyed larvae, the larvae were fixed and the head dissected away from the brain. Next, the forebrain was removed, the midbrain was bisected down the midline, and the two halves were freed from the body by cutting just caudal to the midbrain-hindbrain boundary with a microsurgical knife (Fine Science Tools). Each half of the midbrain was transferred directly into a digestion solution using a capillary pipette with as little PBS as possible. Midbrains were dissected on multiple days, frozen after protease digestion, and then pooled together on the same RNA extraction column. Each extracted RNA sample contained 74 midbrain halves. Total RNA was extracted from the samples using the RecoverAllTM Total Nucleic Acid Isolation Kit for FFPE (Ambion), with the deparaffinization step omitted. RNA was eluted into 60 μl nuclease-free water and its integrity was measured by optical density ratio (OD260/OD280 nm) > 2.0 and RQI > 7.6, where 0 corresponds to fully degraded RNA and 10 corresponds to intact RNA (Experion RNA Highsens Analysis, BioRad). cDNA was synthesized and amplified using the TransPlex® Complete Whole Transcriptome Amplification Kit (Sigma-Aldrich). 144 ng of RNA was used from each sample and all cDNA after amplification had OD260/OD280 nm absorption ratio >1.86. RT-qPCR primers are listed in Table S1 and were designed with a TM of 60°C, using either Primer3 (Invitrogen/ThermoFisher) or QuaniTect primer assays (Qiagen). Primer efficiency was calculated by qRT-PCR standard curve analysis using 1:5 serial dilutions of cDNA over at least four samples. Only primers with an amplification efficiency (AE) of 95-105% were used, as determined by the equation AE = 10^-1/gradient^.

cDNA for qRT-PCR analysis was diluted 1:4 or 1:10 in H_2_O. Samples and standards were run in triplicates on 96 well plates. qRT-PCR amplification was performed using SYBR Green JumpStart Taq ReadyMix (Sigma) or iTaq Universal SYBR Green Supermix (BioRad) on a CFX96 RT qPCR machine (BioRad) for 40 amplification cycles.

The parameters for qPCR reactions examining gene expression levels in left and right sides of the brain consisted of an enzyme activation step of 3 minutes at 94°C followed by cycles of 94°C for 15 seconds, 60°C for 30 seconds and 72°C for 30 seconds. The parameters for qPCR reactions examining gene expression in *lakritz* and wild-type sibling brains consisted of an initial enzyme activation step of 3 minutes at 94°C followed by cycles of 94°C for 15 seconds and 60°C for 30 seconds. When comparing gene expression in left and right sides of intact and denervated brains, fold-change of gene expression was calculated by using the 2^-ΔΔCq^ method using elongation factor 1α (*ef1α*) as a reference gene. When comparing gene expression between wild-type and *lak* mutant brains, due to the dramatic decrease in gene expression, relative normalized expression was calculated as 2^-(ΔΔCq/avΔCqwt)^ for each gene. Melt-curve analysis was used to identify and remove reactions with nonspecific products.

### Image Acquisition and Quantitation

For bright field microscopy, brains were imaged on a Nikon Eclipse E1000 microscope using a Micropublisher 5.0 RTV camera (Q imaging). For laser scanning confocal microscopy, brains were imaged on either a Nikon A1+ laser scanning confocal microscope, using a 25X 1.1 NA long working distance water immersion objective, or a Leica SP8 confocal microscope with 20X 0.8 NA water immersion objective. Resolution was set to 1024x1024 pixels, pixel size was 0.50 µm, and pixel dwell time was 1.1 µs/pixel. Images were taken at 1-2 µm intervals in the Z-plane over a total depth of 100-200 µm, varying by age of larvae and mounting position. For selective plane illumination microscopy, brains were imaged on a MuVi (luxendo/Bruker) at 22X magnification using the 1.6x beam expander and then processed with LuX software. All micrographs were rendered and analyzed using Imaris x64 (Bitplane) or Fiji (ImageJ; (Schindelin et al., 2012)).

To assess differences between the left and right sides of the tectum, image stacks from the confocal microscope were processed with Imaris. A contour surface was created using the DAPI channel to represent the midbrain volume. Using the edit surface tab, the surface was cut at the midline, separating the left and right sides. The two surfaces were used to select the feature to be quantified and normalized for brain size across samples. EdU-positive and TUNEL-positive puncta were observed using the Imaris Spots tool, and spots in the neuropil were manually excluded.

In *lakritz* mutants and their wild-type siblings, two regions of interest, one in the left tectal lobe and one in the right, were drawn in each larval brain at the same position along the anterior-posterior axis and EdU-positive cells within those regions were counted and summed. TUNEL-positive cells were counted throughout the OT with the Spots tool in Imaris and any TUNEL-positive spots in the neuropil were excluded.

To quantify the extent of *sox2* or *wnt3a* expression in the OT, contour surfaces were created for the DAPI channel, and either Fast Red (excited with 561 nm light) or Fast Blue (excited with 640 nm light) channels using Imaris. Due to the extensive amount of *wnt3a* and *sox2* staining, accurately counting individual cells was not possible. Therefore, the volume of the *wnt3a* or *sox2* contour surfaces was calculated relative to the volume of the DAPI-stained volume of either the entire tectum, or the left and right sides of the tectum. When analyzing one-eyed larvae, the contour surfaces were manually cut at the midline using anatomical markers to position the cutline.

To quantify the extent of Wnt-target gene expression in the OT by in situ hybridization, images were converted to black and white and then threshold in Fiji (Schindelin et al., 2012) using an upper limit between 90-110. An ROI of ∼800x360 px was drawn, centered across the midline and the thresholded area was measured. Two ROIs ∼800x180 px (identical size for each image) were then drawn within the first ROI just on either side of the midline, and then the thresholded area in each was measured.

For comparisons of one-eyed versus intact larvae, the staining on the right (denervated) side was subtracted from the left (intact) side and divided by the total staining to generate a difference ratio of (left-right)/total that was plotted and compared. For *lak* mutant and wild-type sibling larvae, total area was plotted and compared.

### Computation and Statistical Analysis

Statistical analyses were performed and plotted using RStudio (Grolemund, 2020; RStudio Team, 2019). A list of packages, libraries, and sample code can be found in supplemental materials (S1 document). Super Plots were adapted from R code provided by (Lord et al., 2020). Datasets were tested for normality with the Shapiro-Wilk test and Q-Q plots, and evaluated for equal variance using Fisher’s F test or the Fligner-Killeen test prior to t-tests, and Levene’s test prior to two-way ANOVAs. Two-way ANOVAs were followed with Tukey’s HSD post-hoc testing. Wilcoxon Ranked Sums tests or Welch’s t-tests were performed for pairwise comparisons and corrected for multiple comparisons. Effect sizes were calculated with Hedge’s g and Omega-squared (ω2) tests. The difference between the means (∂) was calculated by subtracting the average ratios of TUNEL or EdU in tecta of one-eyed larvae from intact larvae. Tables of all summary statistics (Table S2), test statistics (Table S3), and effect size statistics (Table S4) are included as Supplementary Materials.

## Acknowledgements

Thank you to Cerveny and Wilson Lab members past and present who read drafts and/or talked through aspects of this project even though they were working on other (very different) things

## Contributions

Conceived of the initial project - KLC, MV, HR, SWW

Performed experiments, composed figures, analyzed data - KLC, MV, HR, EEK, OH, YK, SH, CR, JY

Wrote and edited the manuscript: KLC, MV, HR, EEK, OLH, YK, CR, SH, JY, SWW

## Supplementary Figures and Figure Legends

**Figure S1:**
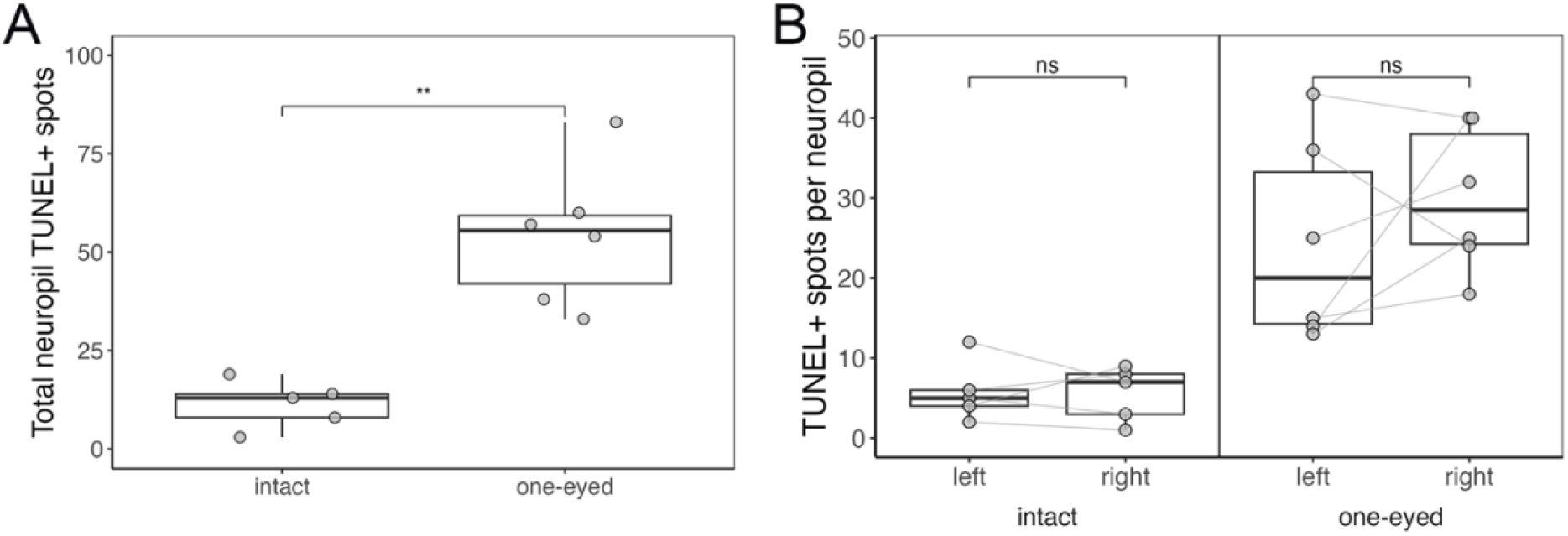
The number of TUNEL+ spots increases in neuropil regions of innervated and denervated sides of the OT one day after eye extirpation. (A) Box plot overlaid with individual data points for total number of TUNEL+ cells in the regions where RGC axons terminate (OT neuropil) for 6 dpf larvae that are either intact or one-eyed (eye removed at 5 dpf) wild-type. (B) Same data as in A, binned into left and right sides. Box plot overlaid with individual data points for number of TUNEL+ puncta on left and rights sides. Dotted lines connect paired samples within individual embryos. For both plots, statistical significance determined by Welch’s two sample t-test; n = 6 for each condition.

**Figure S2:**
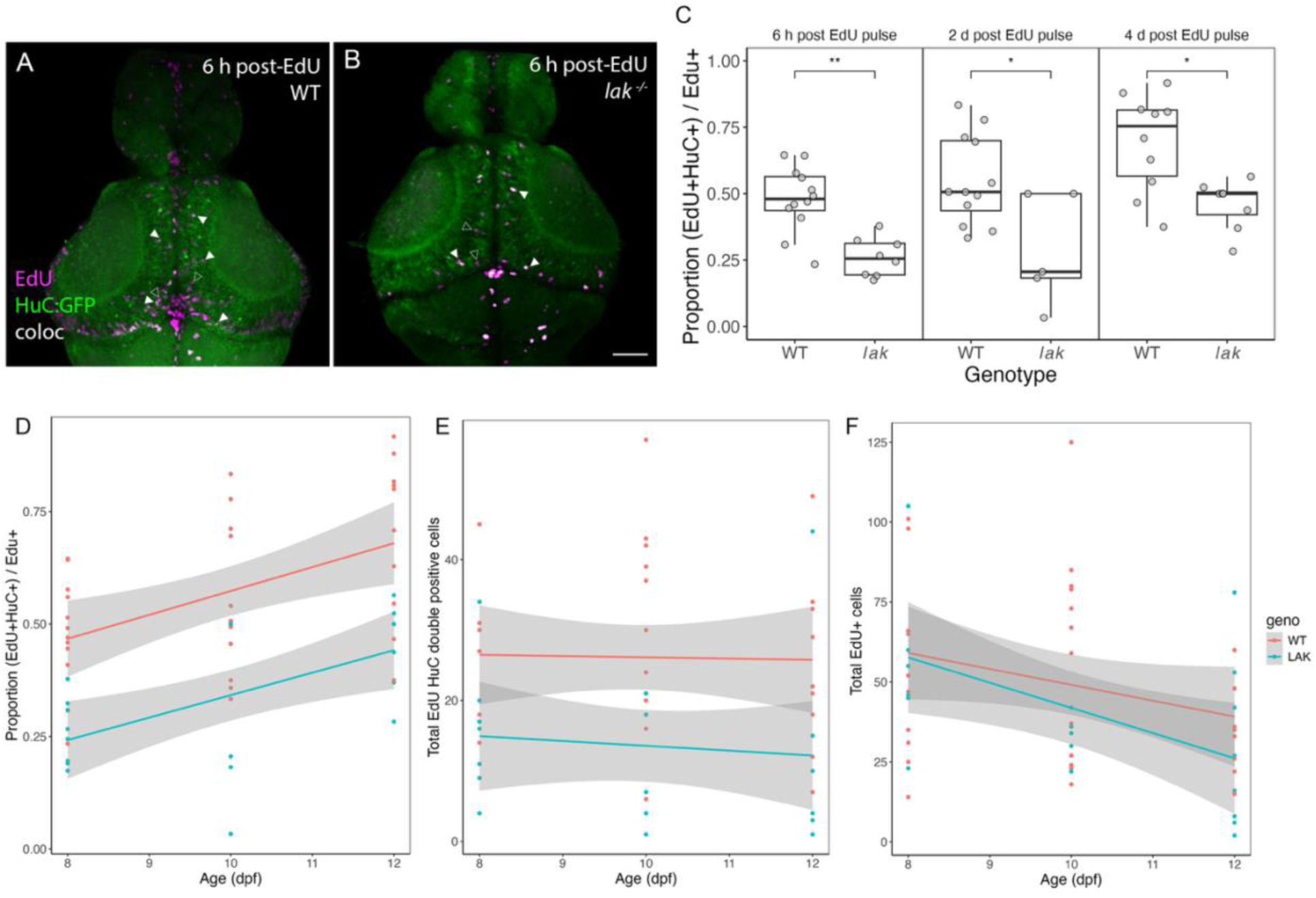
Wild-type and *lak* mutant neural progenitors differentiate into HuC+ neurons at indistinguishable rates. (A-B) Maximum intensity projections of dorsal views of dissected brains from wild-type (A) and *lak^-/-^* (B) larvae with a HuC:GFP transgene to label neurons. Larvae were pulsed with EdU at 8 dpf for 4 h and then washed into embryo medium for 2 h before fixation followed by EdU detection (magenta) and immunostaining for HuC:GFP (green). Colocalization of both EdU+ and HuC+ is visible in white, double-positive cells indicated with filled arrowheads, EdU+ only cells indicated with, open arrowheads. Bar is 50 µm. Ages and genotypes indicated in upper right corner of both panels. **(C)** Box plots overlaid with individual data points show the proportion of double-positive cells relative to total EdU+ cells as a function of genotype and age (8 dpf, 10 dpf, and 12 dpf). n ≥ 5 for each condition. Statistical significance determined by Welch’s t-test. Abbreviations: **** p ≤ 0.00001; ** p ≤ 0.01.**(D-F)** Linear regression models overlaid with individual data points show (D) proportion of EdU+ HuC+ double positive cells relative to total EdU+ cells, (E) total number of EdU+ HuC+ double positive cells, and (F) total number of EdU+ cells for WT (orange) or *lak^-/-^* (blue) larvae at times: 6 h, 2 d, and 4 d after EdU pulse. 95% confidence interval shaded in grey.

**Figure S3:**
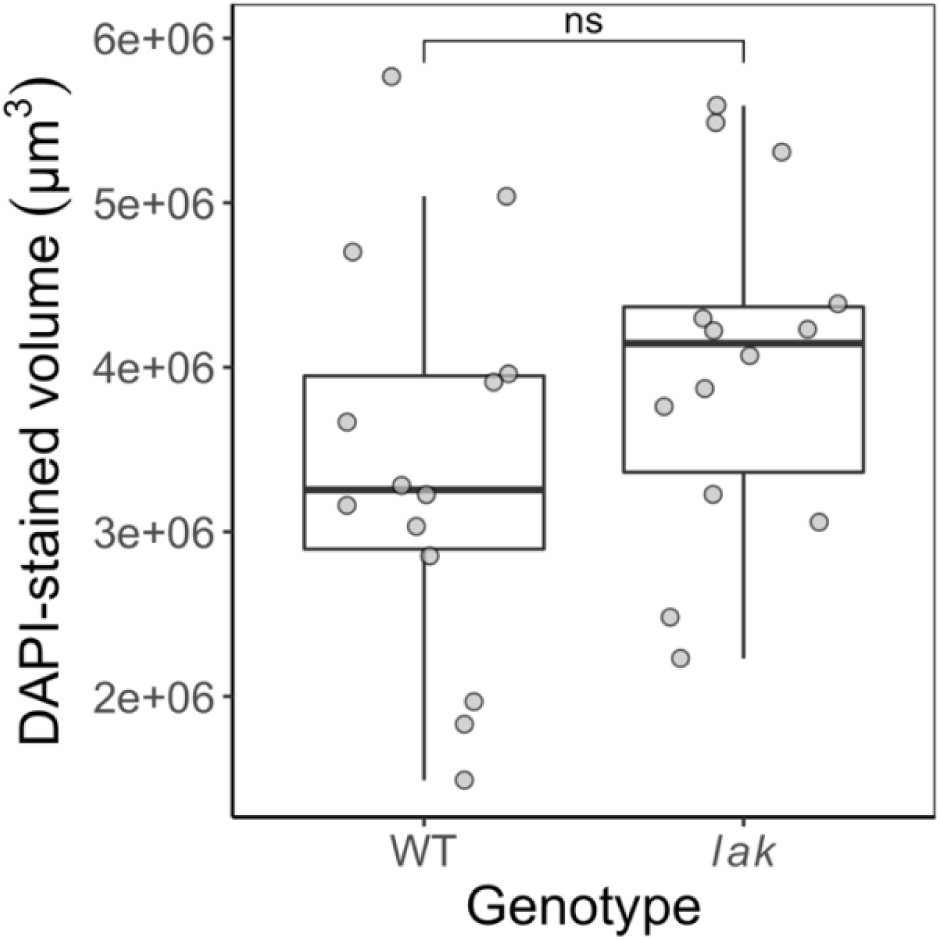
Wild-type and *lak* mutant OT volumes are similar at 9 dpf. Box plot overlaid with individual data points showing the volume (µm^3^) of the DAPI-stained region of the optic tectum for wild-type and *lak^-/-^*larvae at 9 dpf. Statistical significance determined by Welch’s two sample t-test; n=14 for both genotypes.

**Figure S4:**
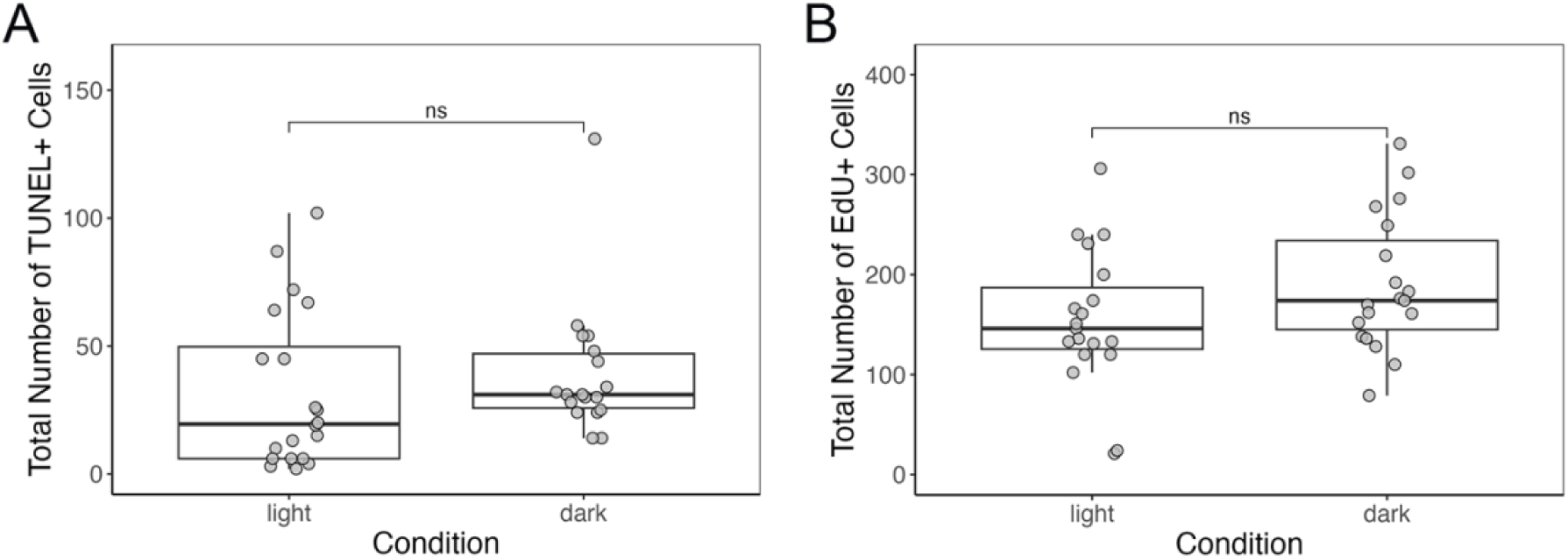
Total number of proliferating or dying cells in the 9 dpf OT appears unchanged in light-dark or constant-dark environments. (A) Box plot overlaid with individual data points showing the total number of TUNEL+ cells for intact wild-type larvae reared until 9 dpf in the indicated conditions. (B) Box plot overlaid with individual data points showing the total number of EdU+ cells for intact wild-type larvae reared until 9 dpf in the indicated conditions. Statistical significance determined by Welch’s two sample t-test; n = 18-20 for each condition.

**Figure S5:**
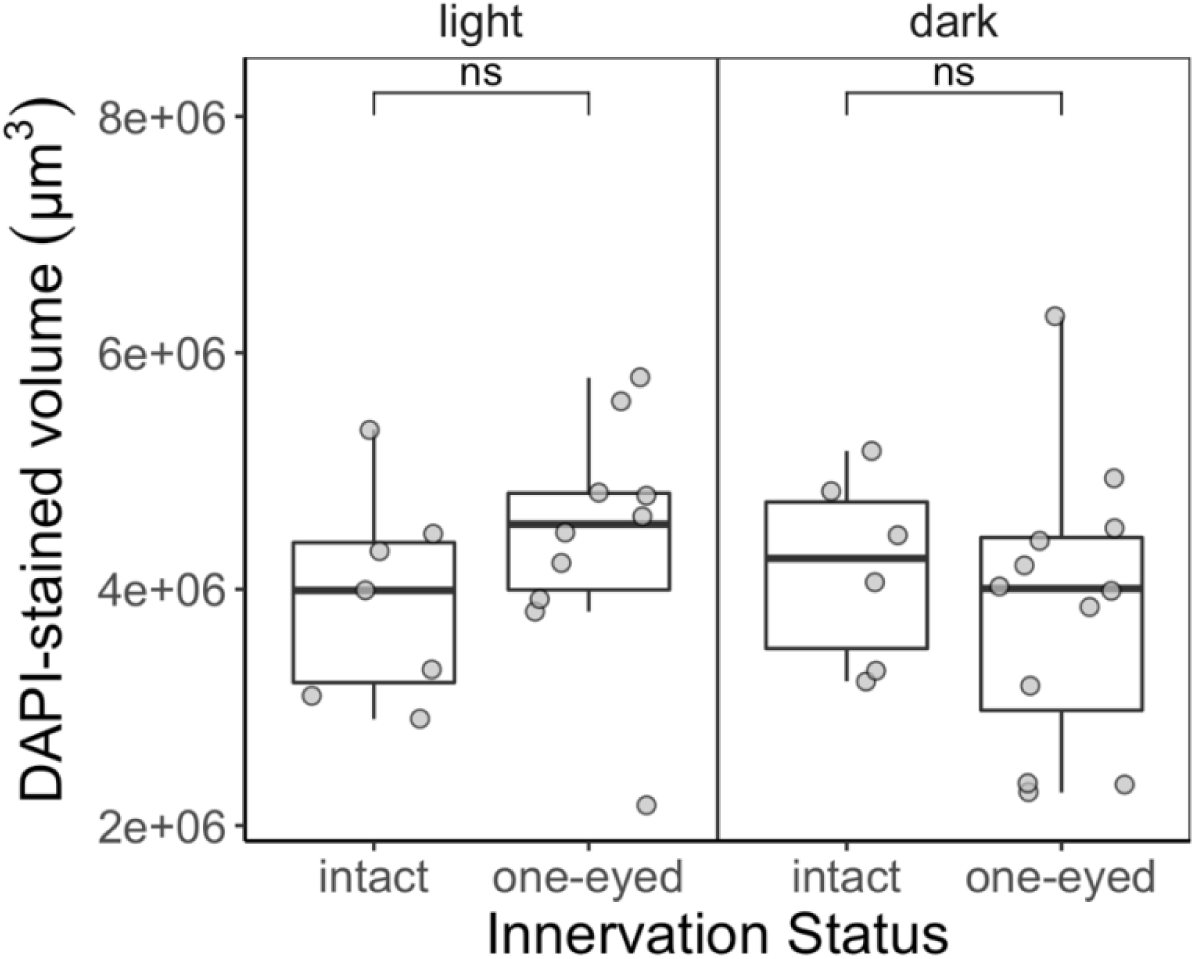
Dark rearing and eye extirpation have minimal effects on optic tectum volume. Box plot overlaid with individual data points showing the volume (µm^3^) of the DAPI-stained region of the optic tectum for one-eyed and intact wild-type larvae reared until 9 dpf in the indicated conditions. Abbreviations: ns, not significant. Statistical significance was determined by ANOVA with Tukey HSD for multiple comparisons; n ≥ 6 for each condition.

**Figure S6:**
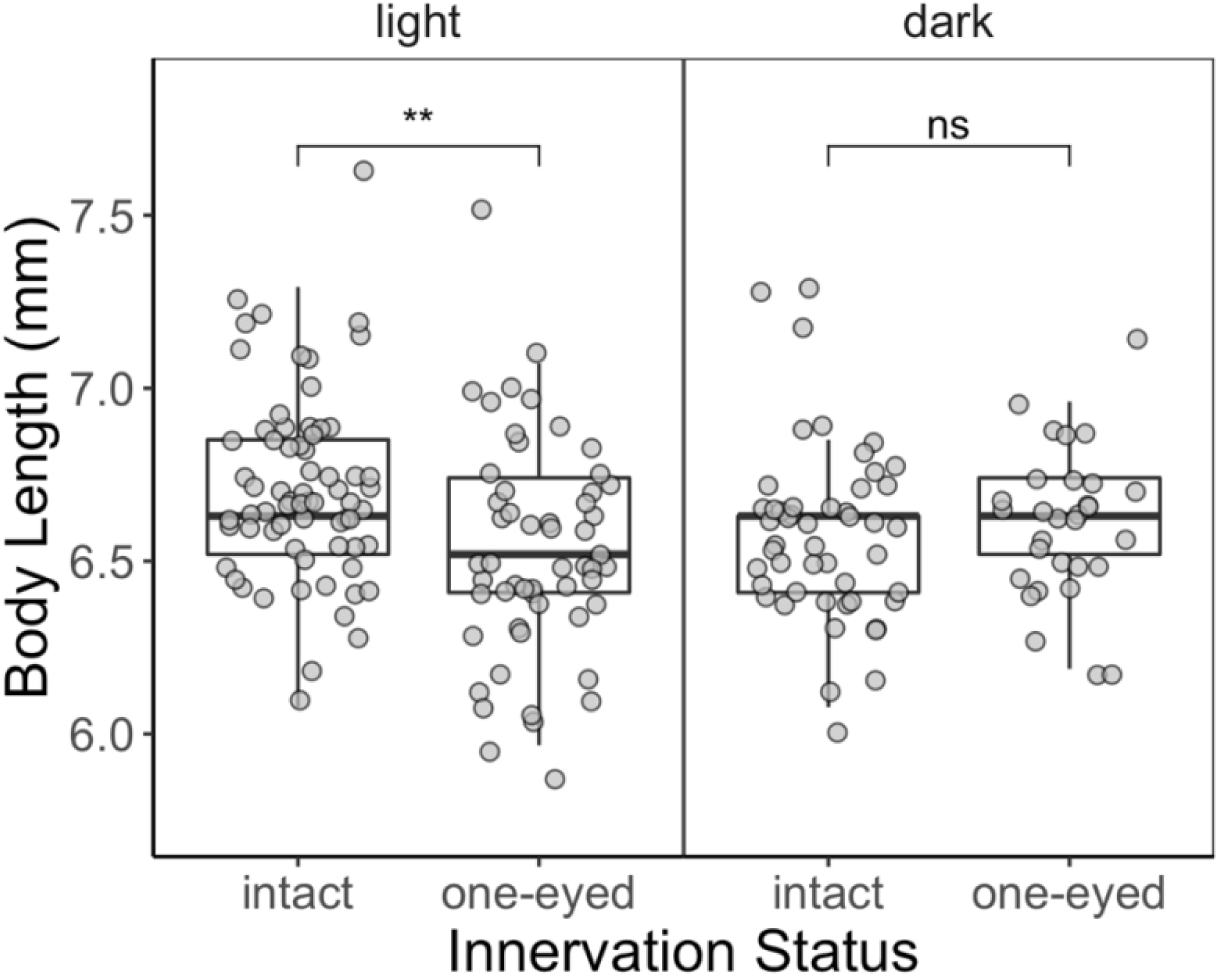
**Dark rearing and eye extirpation have minimal effects on body length.**Box plot overlaid with individual data points showing the body length (mm) from nose to tail tip for one-eyed and intact wild-type larvae reared until 9 dpf in the indicated conditions. Abbreviations: ns, not significant; ** p ≤ 0.01. Statistical significance determined by ANOVA with Tukey HSD for multiple comparisons; n ≥ 30 for each condition.

**Figure S7:**
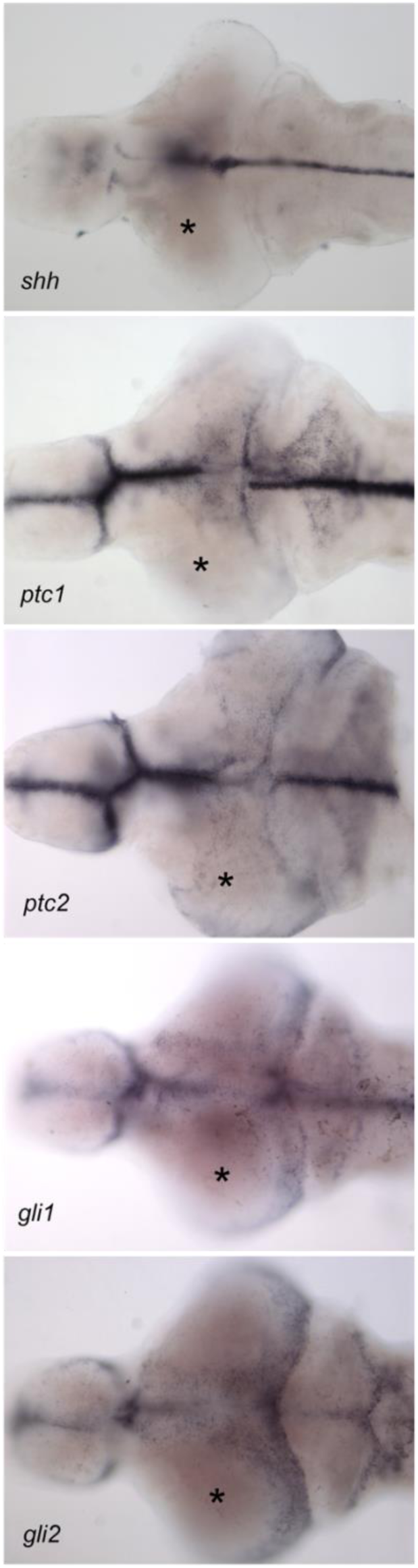
Shh and Hh pathway genes are expressed similarly in innervated and denervated tecta of wild-type larvae. Dorsal views of dissected brains from 9 dpf one-eyed larvae showing expression of mRNA transcripts of the genes labeled in the lower left of each panel by in situ hybridization. Innervated tectal lobes indicated with an asterisk.

**Table 1:**
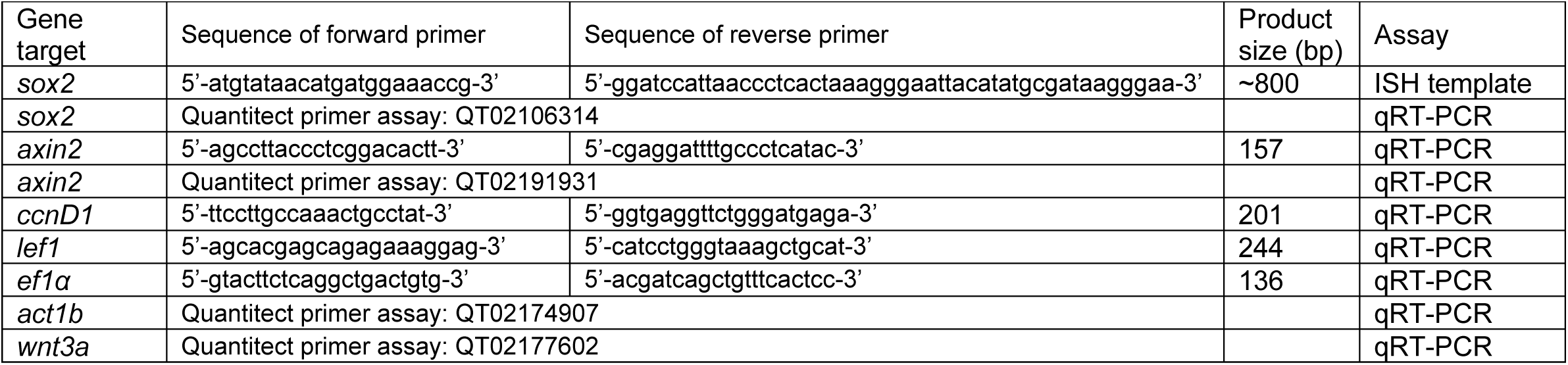
Oligonucleoties.

---

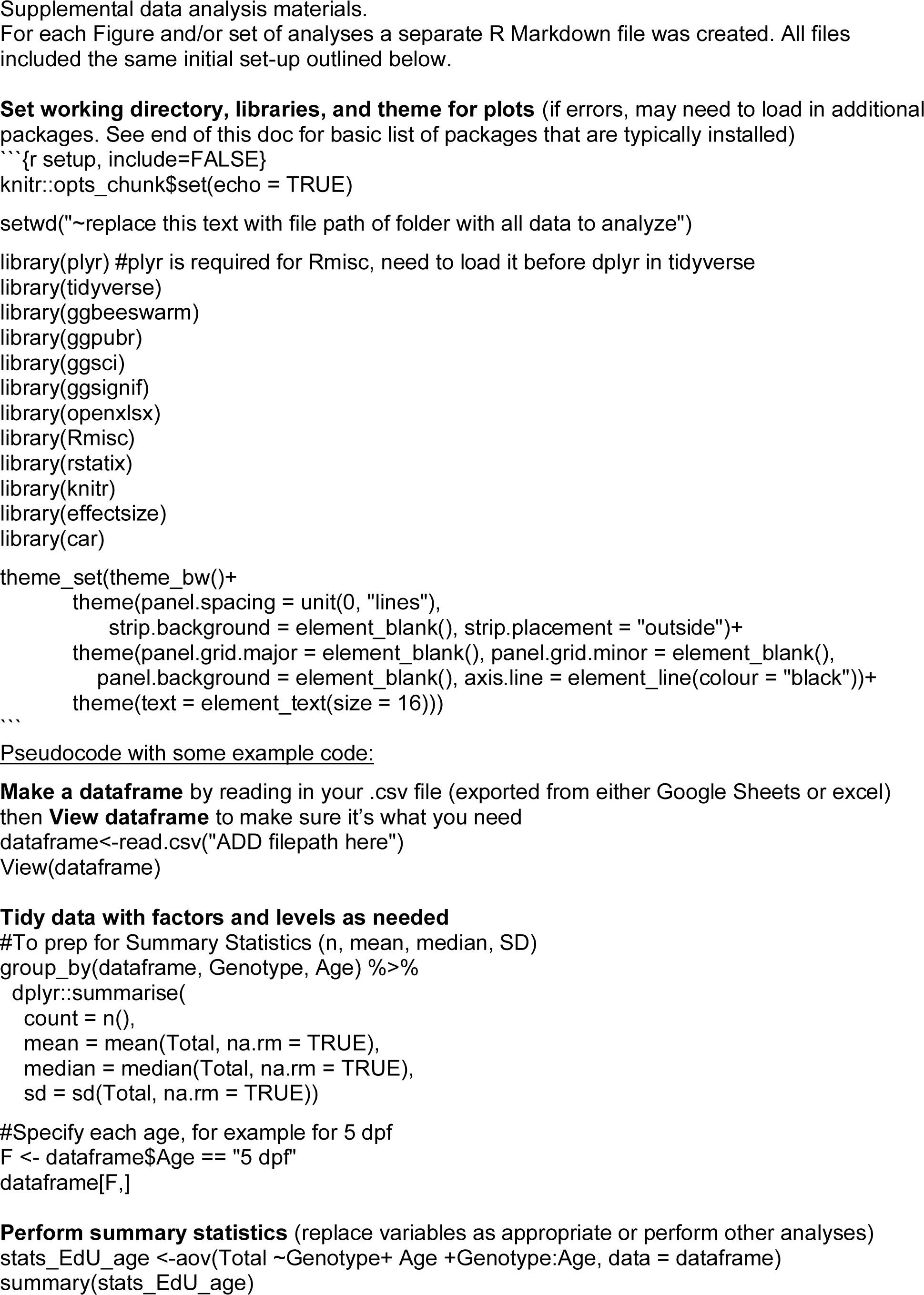
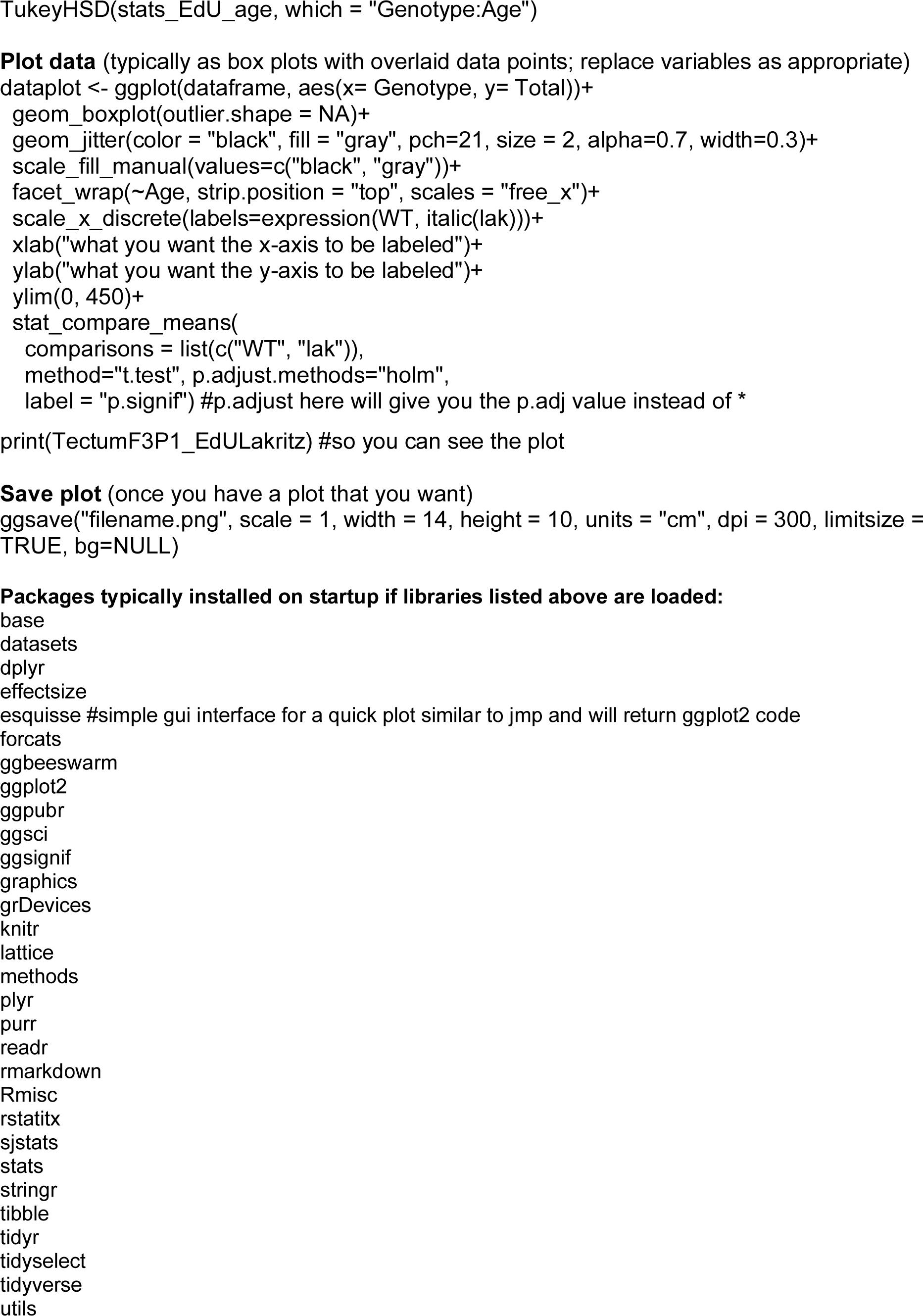

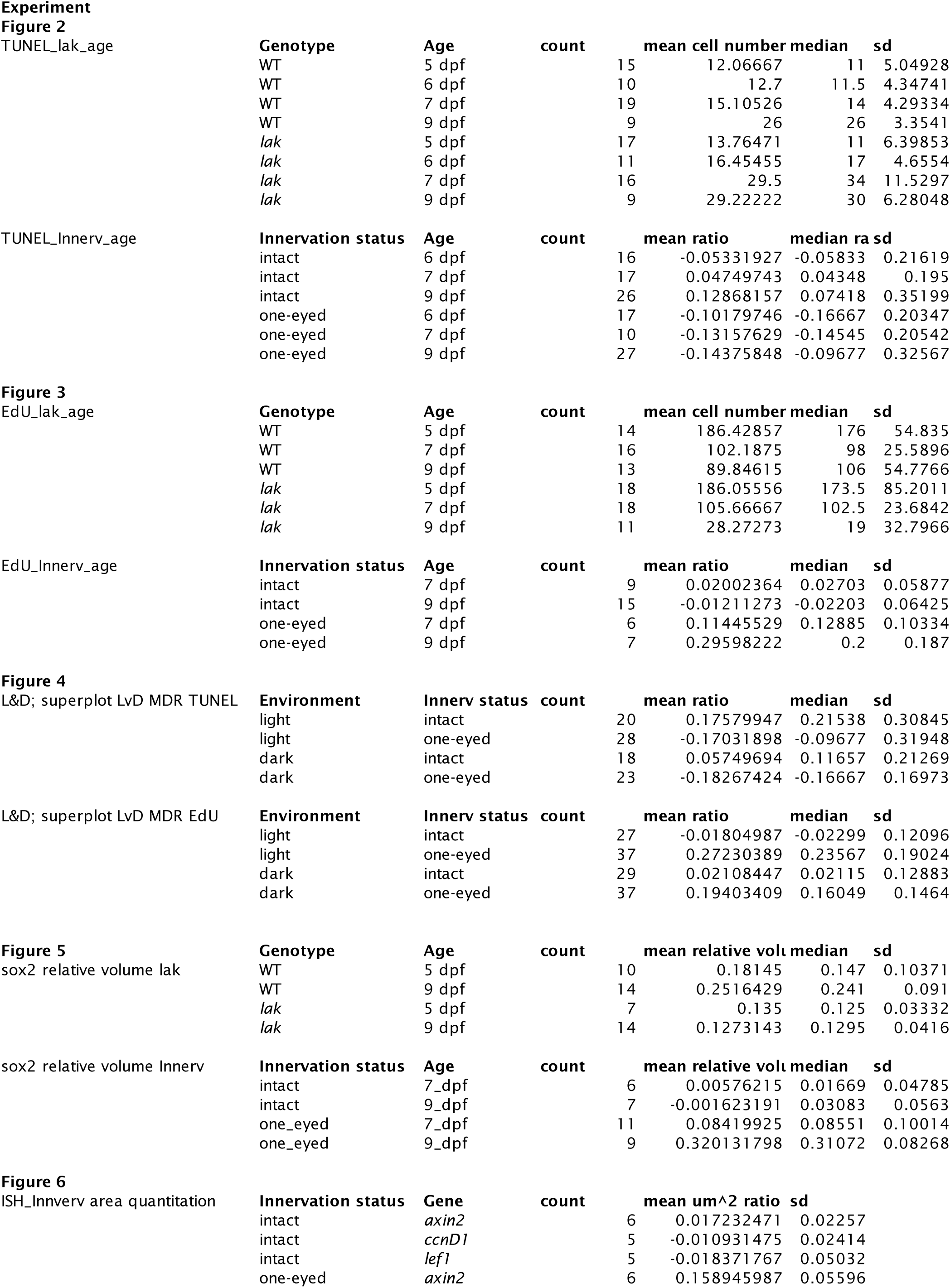
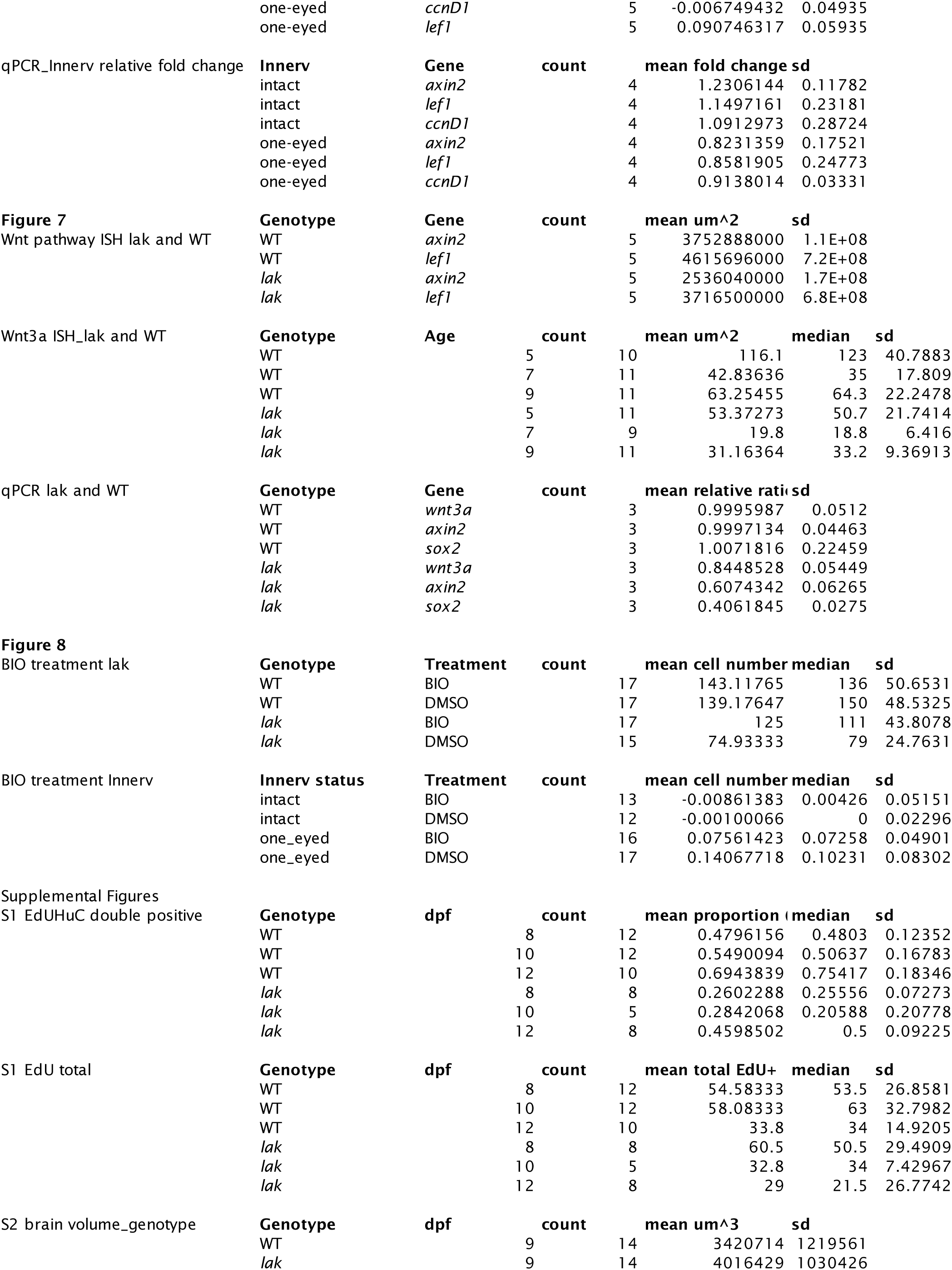
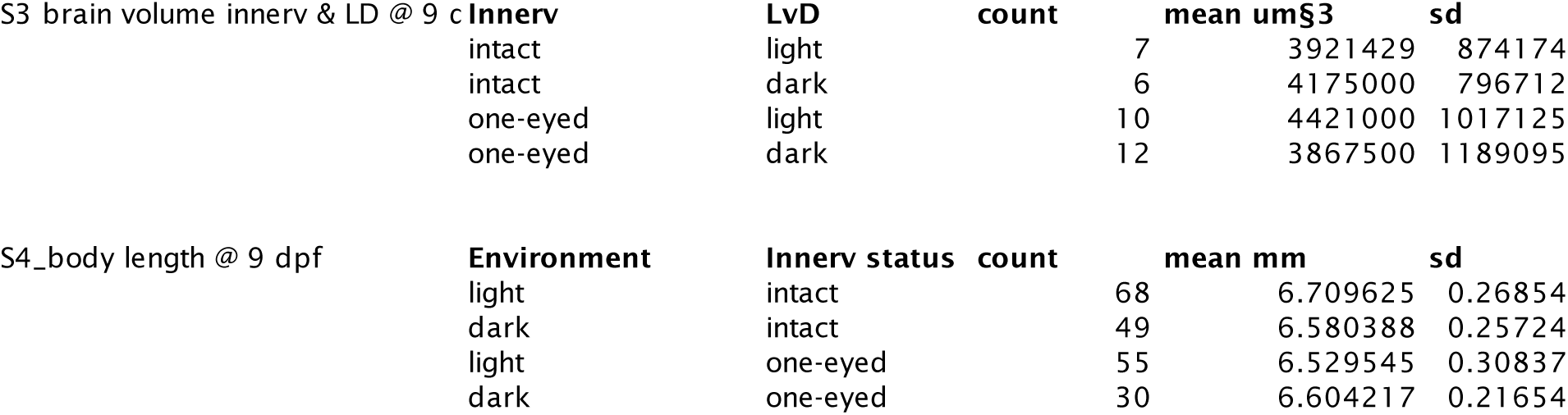
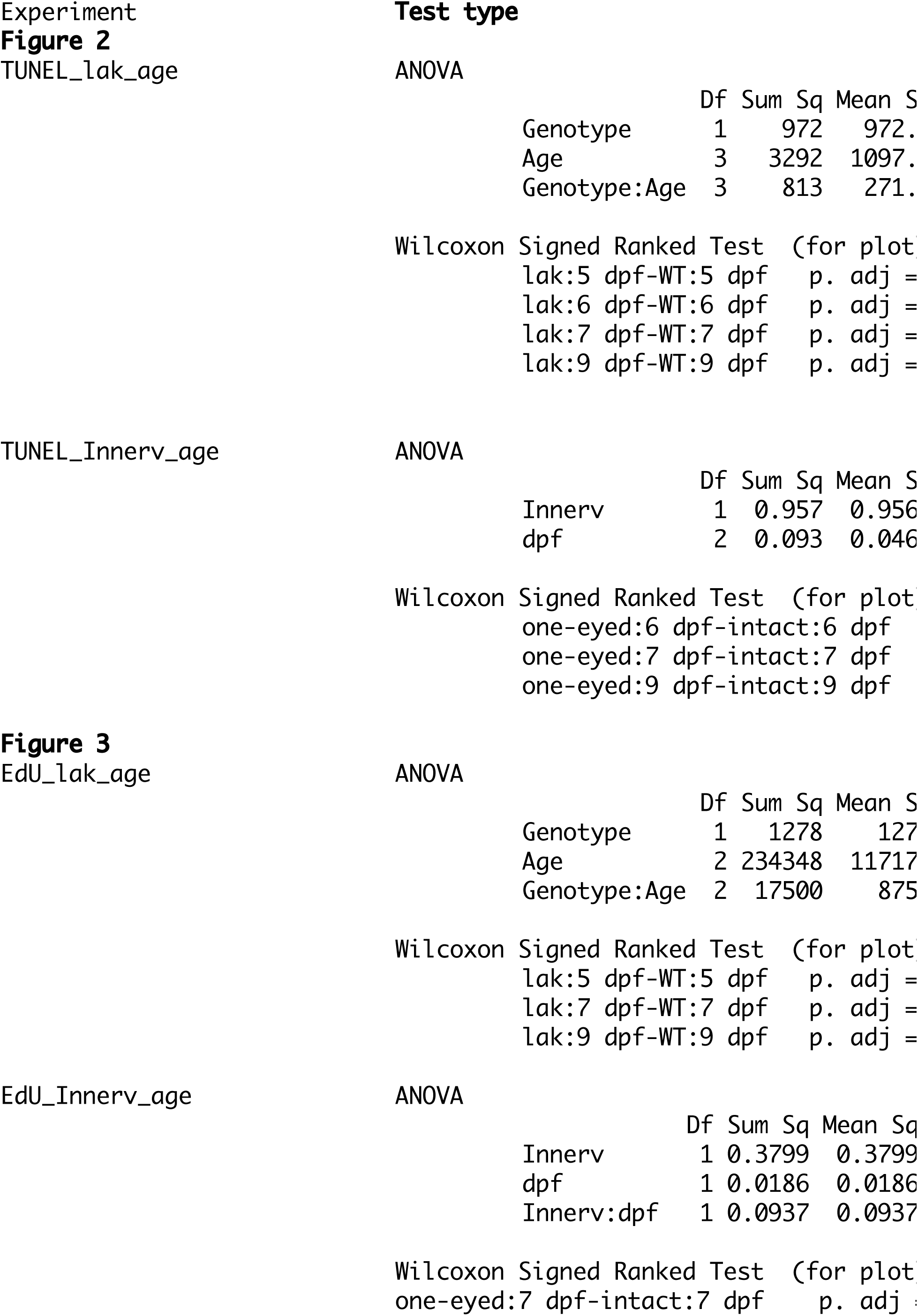
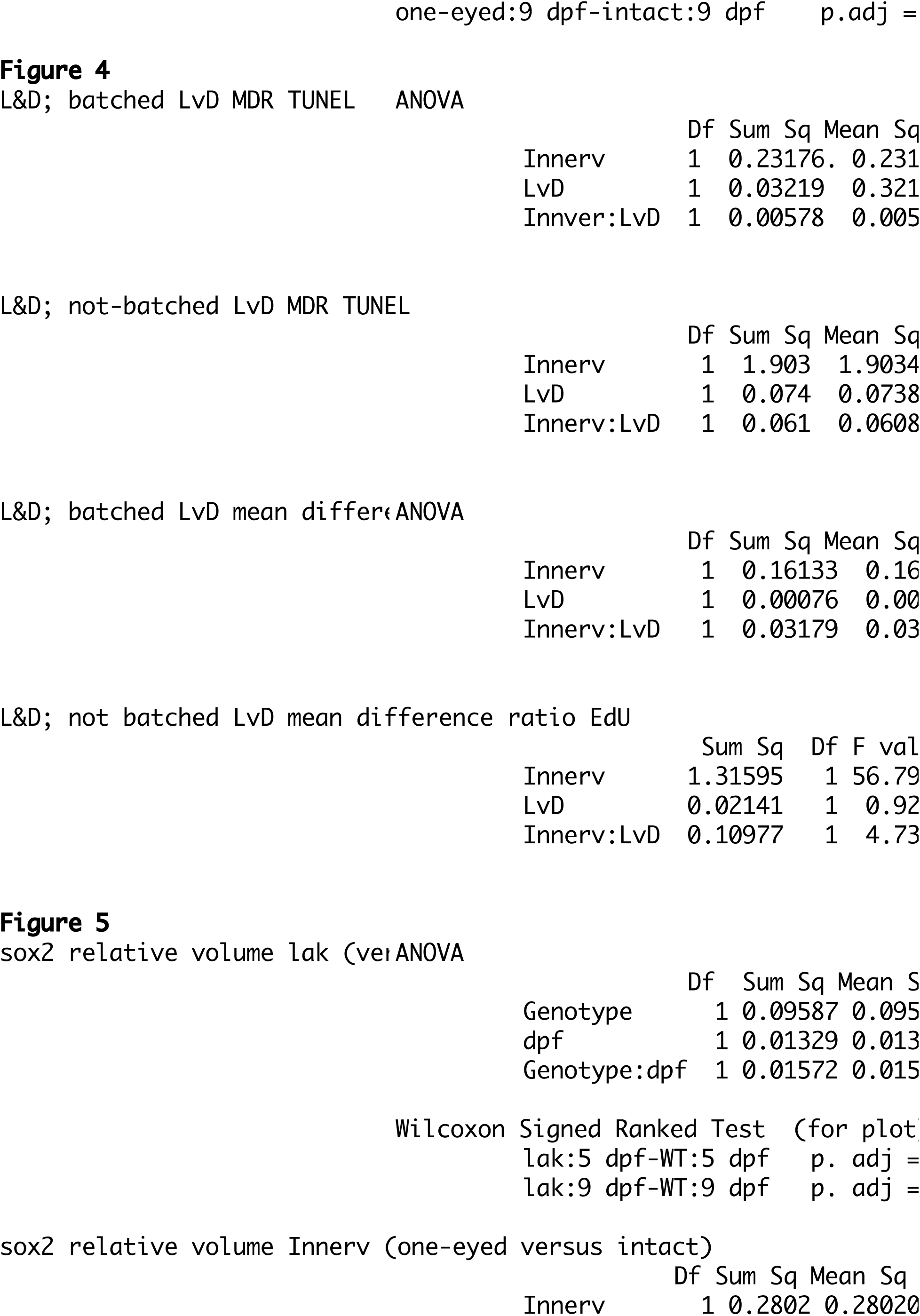
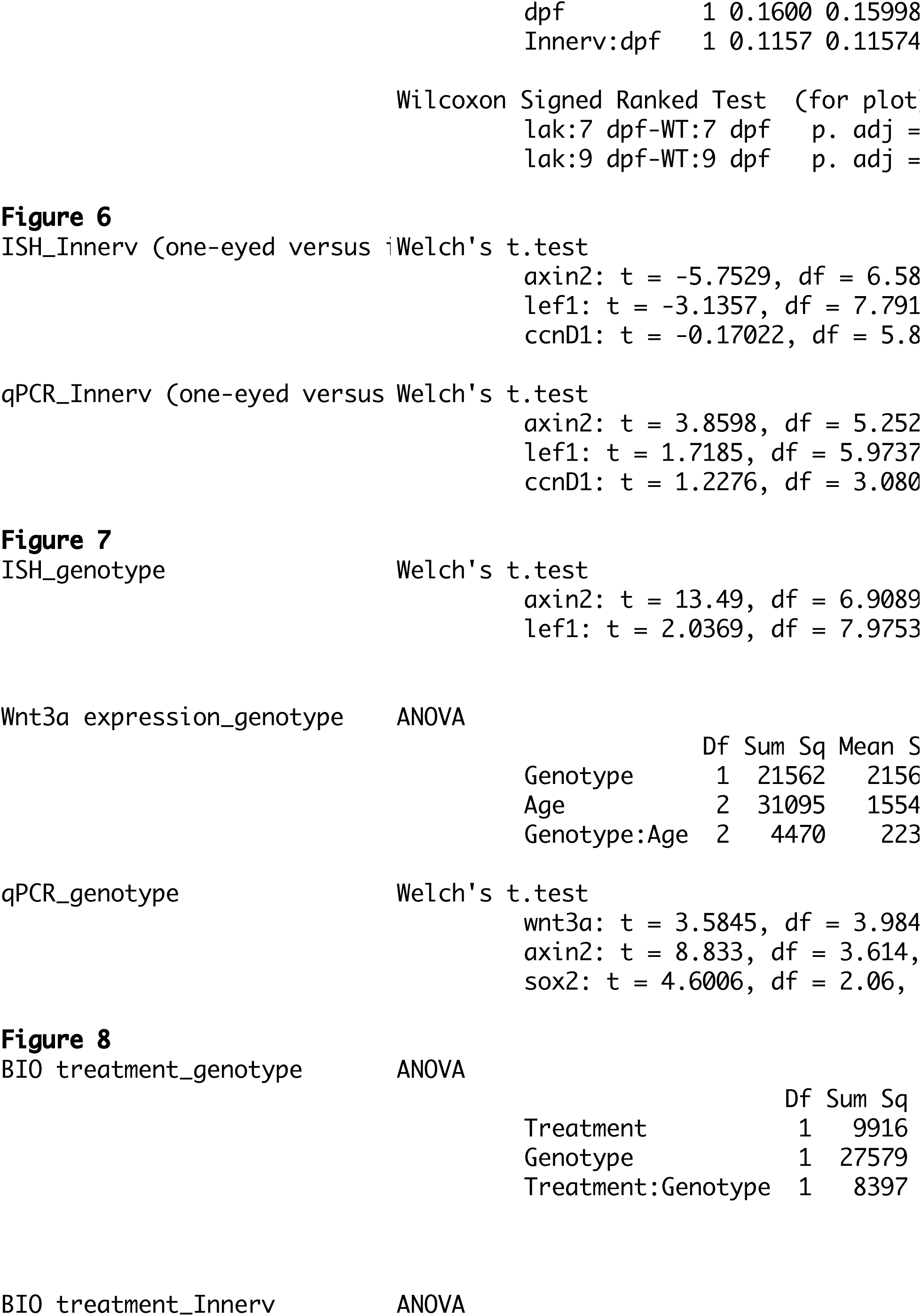
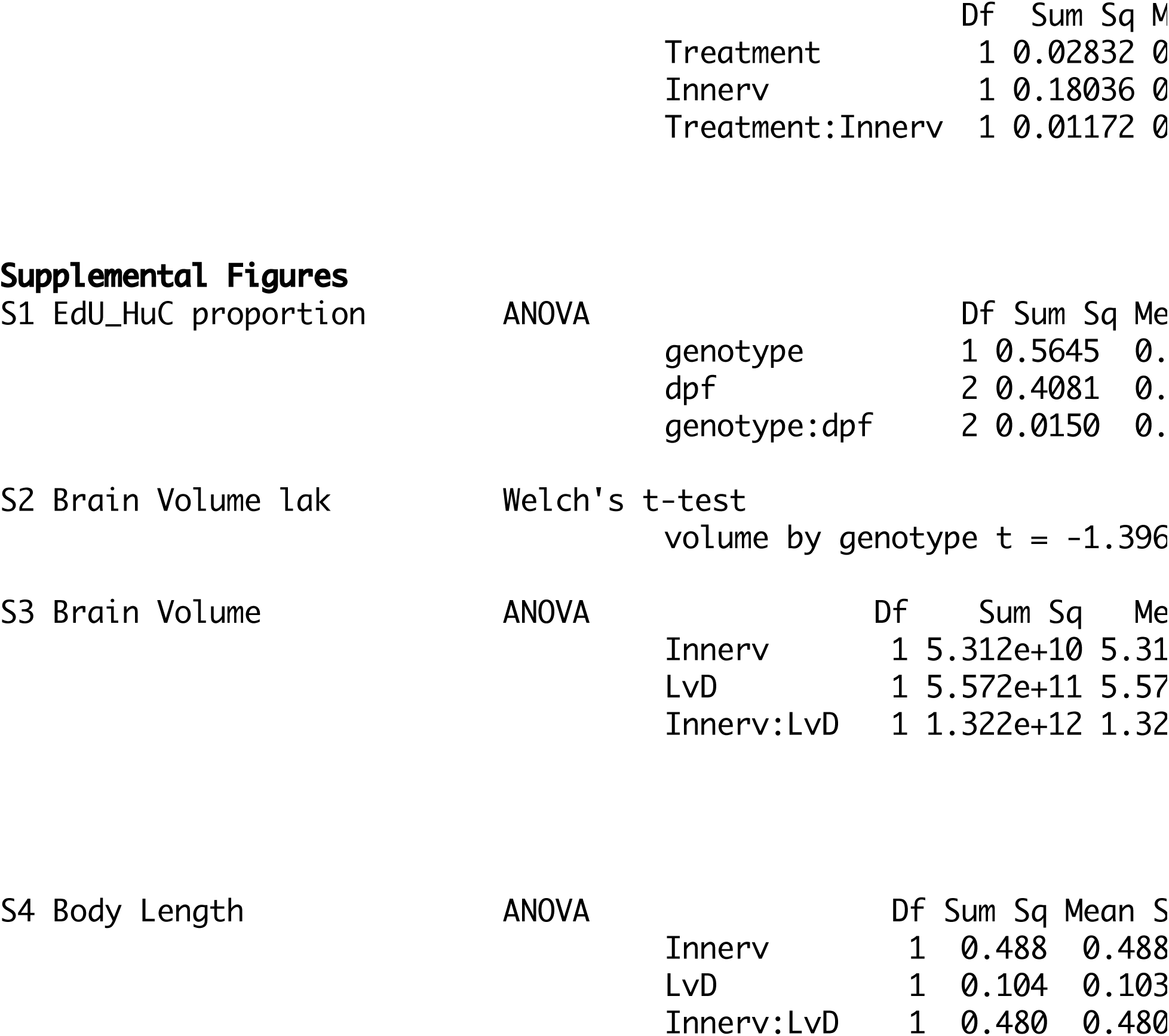
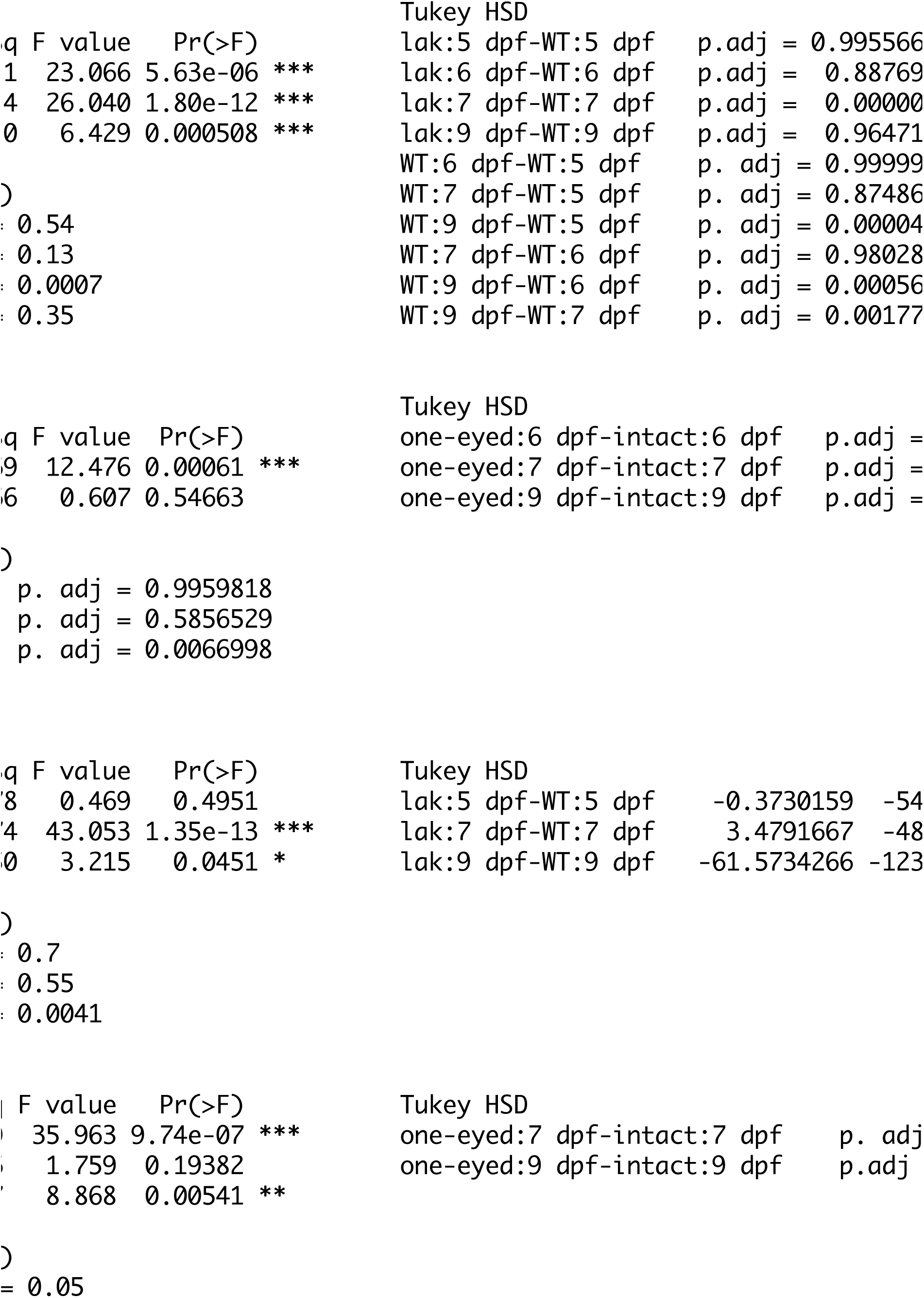
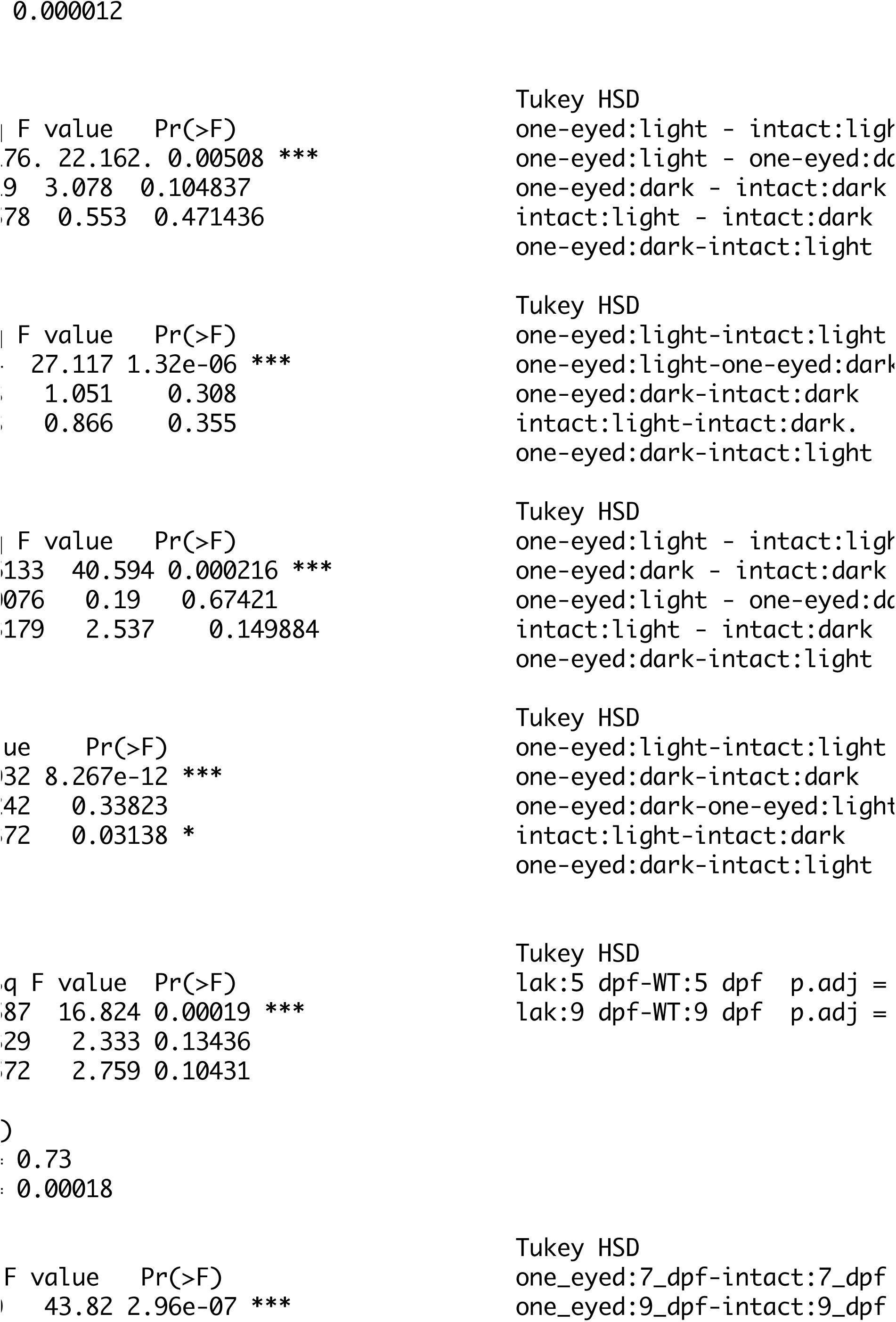
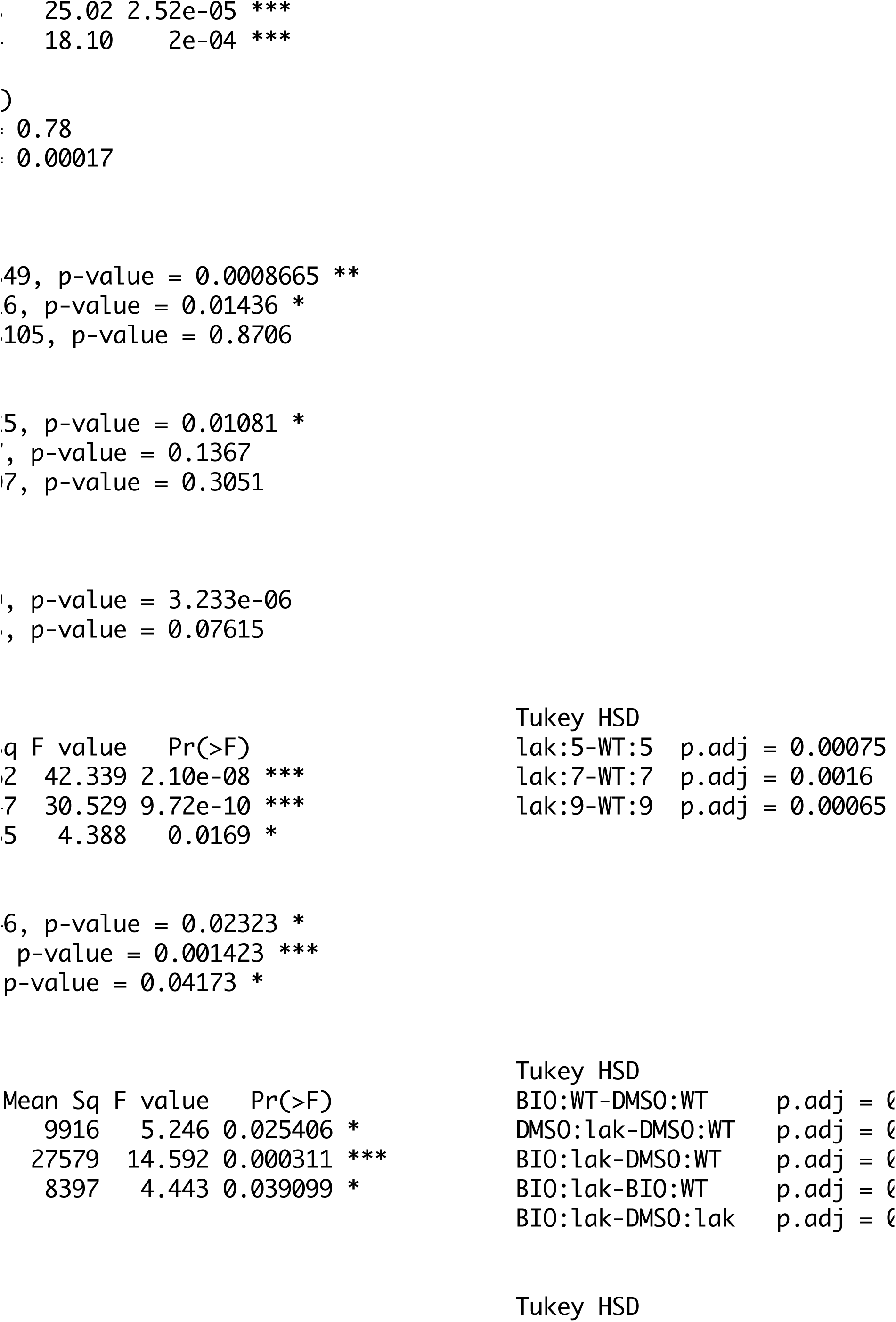
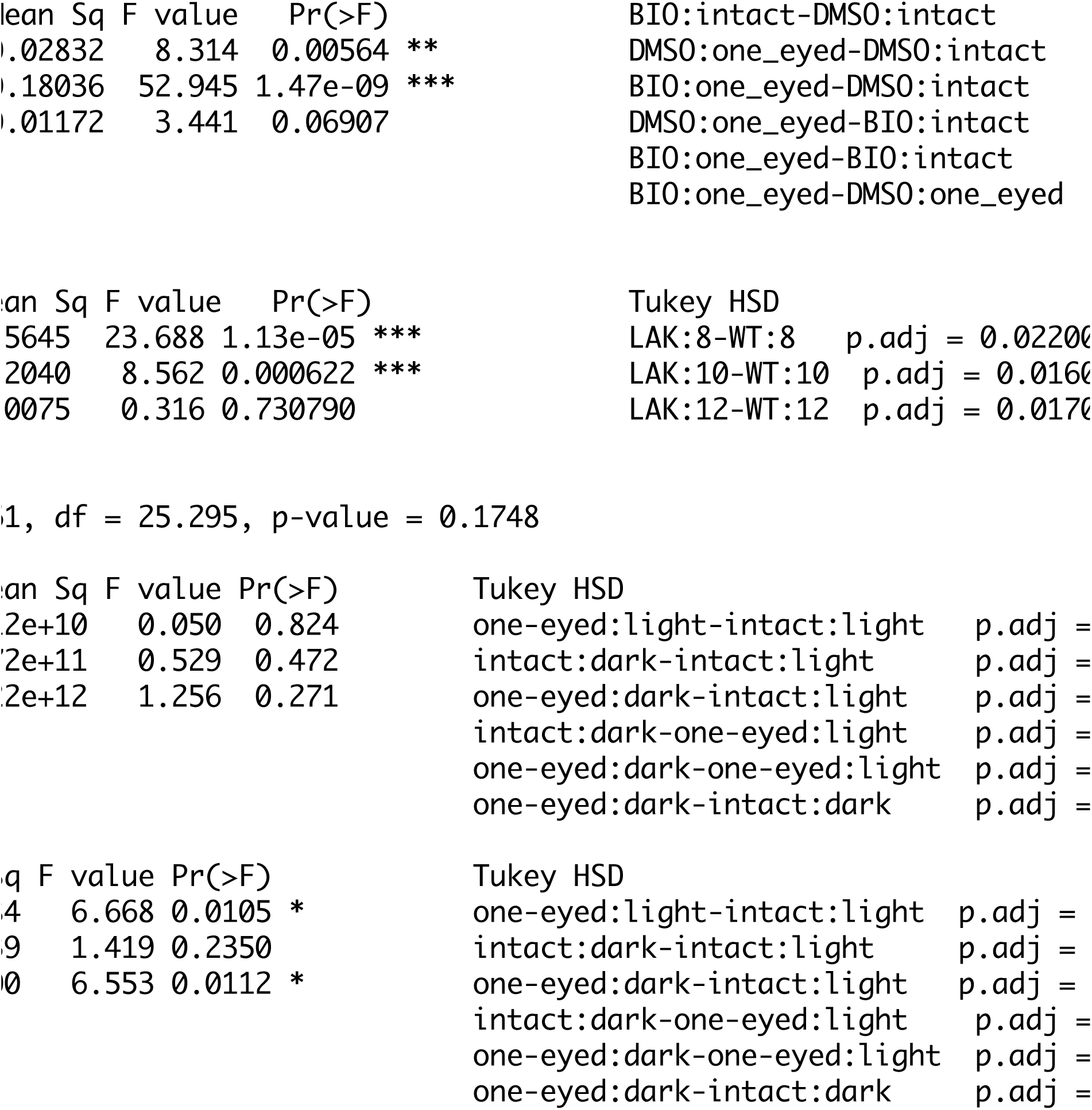
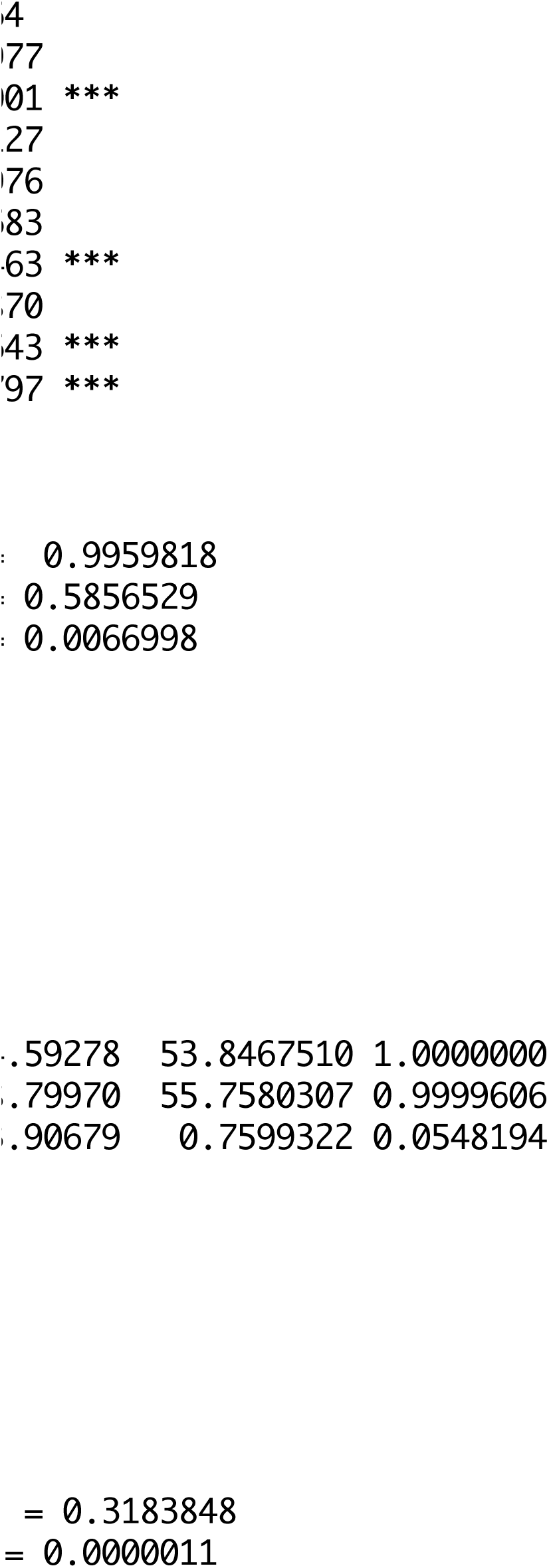
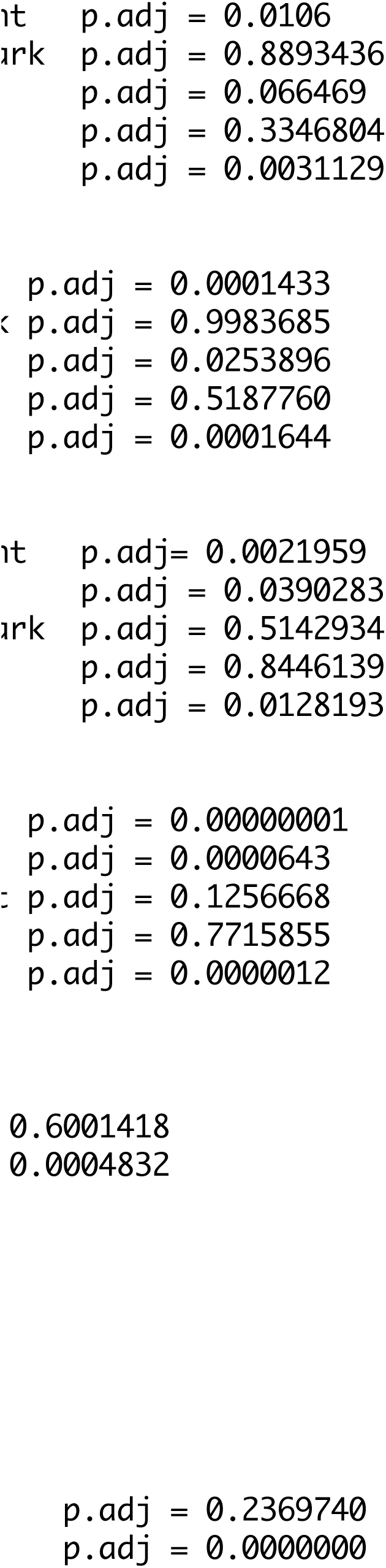
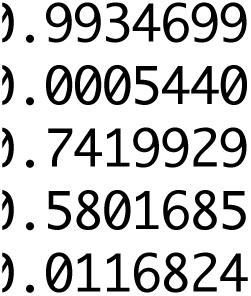
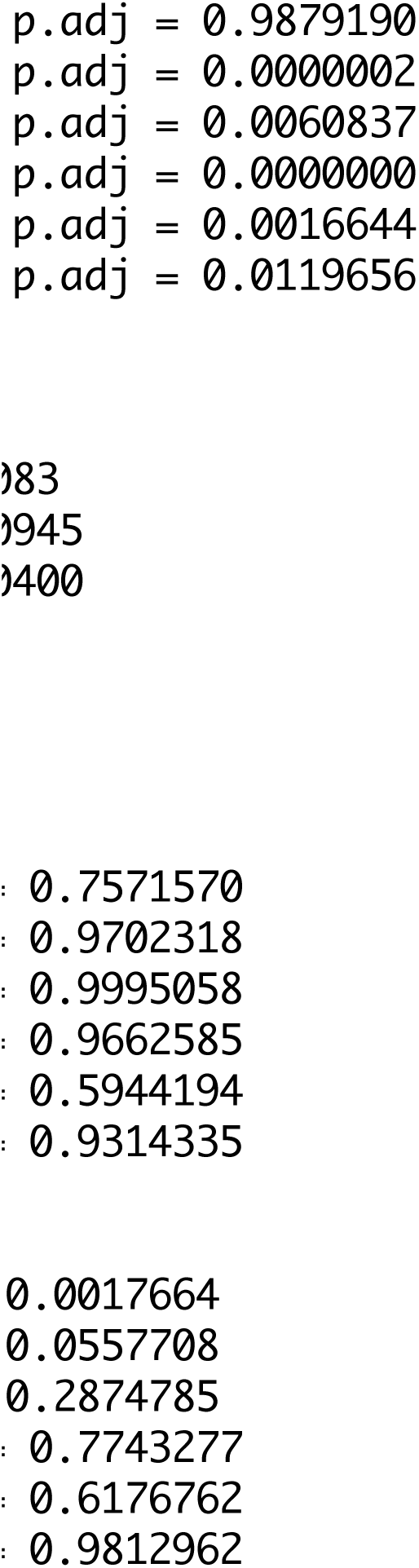
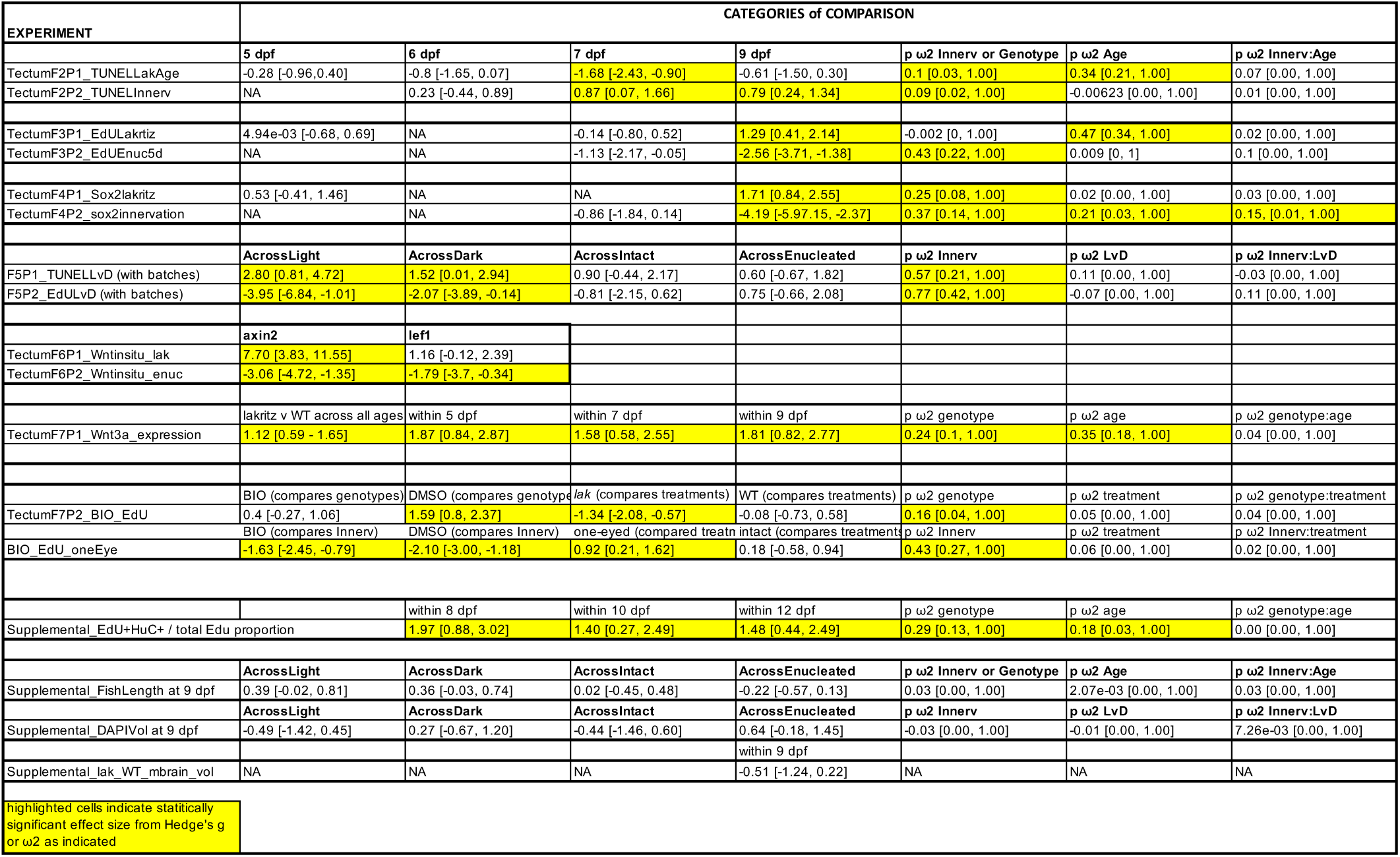

